# EASI-ORC: A Pipeline for the Efficient Analysis and Segmentation of smFISH Images for Organelle-RNA Colocalization Measurements in Yeast

**DOI:** 10.1101/2024.10.04.616601

**Authors:** Shahar Garin, Liav Levavi, Jeffrey E. Gerst

## Abstract

Analysis of single-molecule fluorescent *in situ* hybridization (smFISH) images is an arduous and time-consuming task that is important to perform accurately in order to translate image data into a quantifiable format. This task becomes increasingly more difficult greater the experimental scope and number of images captured. Although smFISH is the gold standard for RNA localization measurements, there are no freely available, user-friendly applications for assaying messenger RNA localization to sub-cellular structures, like the endoplasmic reticulum (ER) or mitochondria). We have developed a pipeline that allows for the automated analysis of multiple smFISH images in yeast cells: EASI-ORC (<u>E</u>fficient <u>A</u>nalysis and <u>S</u>egmentation of smFISH Images for <u>O</u>rganelle-<u>R</u>NA <u>C</u>olocalization). The EASI-ORC pipeline automates the segmentation of cells and sub-cellular structures (*e.g.* organelles), identifies *bona fide* smFISH signals, and measures the level of colocalization between an organelle and mRNA signals. Importantly, EASI-ORC works in a fast, accurate, and unbiased manner that is difficult to replicate manually. It also allows for the visualization of data filtering and outputs graphical representations of the colocalization data along with statistical analysis. EASI-ORC is based on existing ImageJ plugins and original scripts, thus, allowing free access and a relative ease of use. To circumvent any technical literacy issues, a step-by-step user guide is provided. EASI-ORC offers a robust solution for smFISH image analysis for both new and experienced researchers - one that saves time and effort, as well as providing more consistent overall measurements of RNA-organelle colocalization in yeast.

## Introduction

The sub-cellular distribution of messenger RNA (mRNA) is a basic phenomenon of all life forms^1–5^. Specifically, mRNA localization to different organelles or other sub-cellular compartments has been shown to have a significant effect on cellular functions, likely by allowing for the localized translation of proteins required for these functions. mRNAs have been found to localize to centrosomes^6,7^, different neuronal sites^8–11^, the nuclear envelope^6^, as well as organelles like the endoplasmic reticulum (ER)^12–15^, mitochondria^16,17^, and peroxisomes^18^.

The intracellular localization of RNA is studied using many different approaches. Since its inception^19–21^, smFISH remains the gold standard method for research of sub-cellular RNA localization^22^. Regardless of the smFISH method being used, any analysis of the produced images must be performed accurately and consistently to translate the data into a quantifiable format. Naturally, with an increase in scale of a given experiment, more images are produced and more data is collected. Thus, the task becomes more challenging to perform and can become exceedingly arduous and time-consuming. That is particularly true for organelle-RNA colocalization assays, which involve comparisons between different types of signals and physical structures. The act of judging whether the location of a specific smFISH signal corresponds with that of an organelle is crucial. Moreover, when it is performed manually the judgment can be subjective and error prone.

As such, automation of image analysis is beneficial. It can remove barriers to experiments of larger scope by increasing the rate at which images are analyzed. Also, use of a fixed set of rules can provide much-needed consistency in the colocalization measurements. It can also yield a higher amount of data from each image, by identifying signals that may be unclear to the human eye. Many computational solutions were developed to specifically assist with smFISH signal identification (*e.g*. FISH-quant v2^23^, RS-FISH^24^, deepBLINK^25^, and DypFISH^26^, to name a few. They all provide robust tools for smFISH spot identification and allow for a certain level of automation if the user has the appropriate programming knowledge. However, no such tool exists to jointly facilitate RNA and organelle colocalization. Thus, one that measures RNA-organelle colocalization by fluorescence imaging would be beneficial, especially for users lacking programing knowledge.

The basic challenge of using computational image analysis is the translation of an image into a form where all areas of interest are well-defined. For example, there can be no ambiguity whether a specific pixel encompasses an organelle or a FISH signal, or whether it is inside or outside of the cell. To overcome the lack of available software to perform accurate mRNA-organelle colocalization measurements using smFISH with yeast we used several freely available methods and automated them via our original scripts to create the <u>E</u>fficient <u>A</u>nalysis and <u>S</u>egmentation of smFISH Images for <u>O</u>rganelle-<u>R</u>NA <u>C</u>olocalization (EASI-ORC) pipeline. EASI-ORC performs the segmentation of yeast cells as well as sub-cellular structures, identifies smFISH spots, and resolves RNA-organelle colocalization (Figure 1A-E). Each module of EASI-ORC is used to segment an image for one type of signal (*e.g.* cell, organelle, mRNA). Once an entire image is segmented and each pixel resolved as having a specific type of signal, colocalization can be measured accurately and consistently. As the basis for this pipeline, we elected to use the scripting feature of the ImageJ platform to make sure EASI-ORC can be used freely and provide a step-by-step user guide for the different scripts so that it can be used by any researcher, regardless of their technical proficiency (Figure 1G). A statistical analysis tool was also developed to help analyze and visualize the data collected using EASI-ORC (Figure 1F).

**Figure 1.**
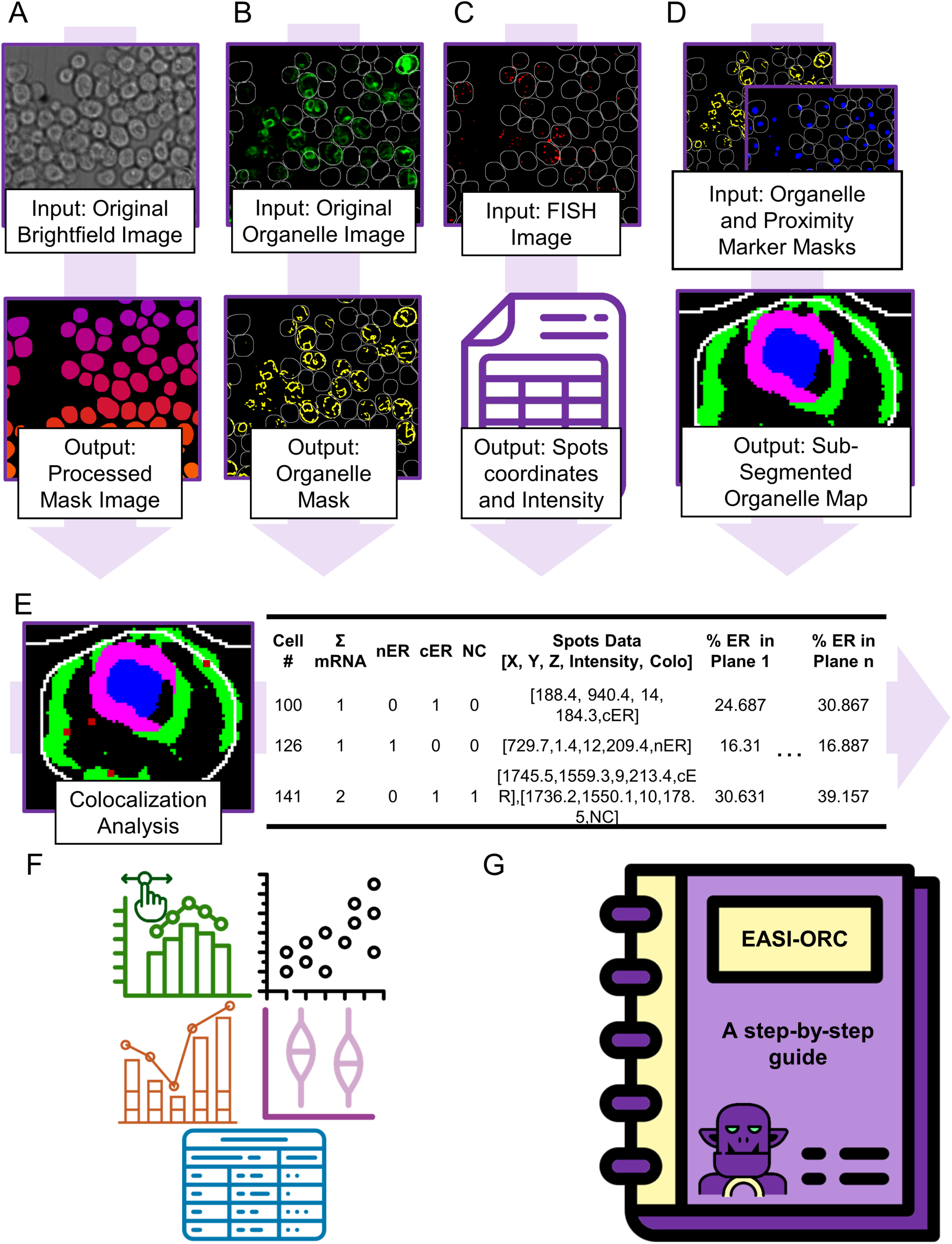
The EASI-ORC pipeline and modules. EASI-ORC modules are designed to work sequentially, but each module can be used independently. **A)** Yeast cell segmentation performed using YeastMate. Shown above, a brightfield image of yeast, while underneath the output mask image is represented by an image marking each identified cell with a discrete color. **B)** Sub-cellular segmentation performed by trainable WEKA segmentation and a user-generated model. Shown above is an image of yeast with fluorescently-labeled ER (via Sec63-GFP) in green. Shown underneath is the output mask image shown in yellow. A white outline marks the identified cells’ boundaries. **C)** smFISH spot identification performed with RS-FISH. Shown above is an image of the signals from the smFISH probe set (*e.g. SUC2* probes labeled by TAMRA). The white outline marks the identified cells’ boundaries. The output is a table with the data of each spot (*i.e.* coordinates and intensity). **D)** Sub-segmentation of organelles performed using proximity marker masks. Exemplified using nER and cER sub-segmentation from the whole organelle and nuclear masks (above, shown in yellow and blue respectfully). Output map represents a single cell example with cER in green, nER in magenta, and nucleus in blue. **E)** Colocalization analysis exemplified using the output mask files and smFISH spots identified in a single *z*-plane. The organelle mask (ER) is represented in green for cER and magenta for nER. smFISH spots are shown in red. White outlines delineate the cell boundary. The output table contains data for every cell identified (each represented as a single row), including the number of spots for each designation (NC = not colocalized), the data for each spot, and the organelle coverage of the cell in each plane. **F)** Toolsets for data filtering, analysis, and visualization. Filtering by smFISH signal intensity, organelle coverage per plane, and number of smFISH signals per cell is dynamically updated. Graphs illustrate RNA spot colocalization with an organelle, the average number of smFISH signals per cell, and important correlations. A statistical summary table is outputted with relevant t-tests and Pearson correlation data. **G)** A step-by-step user guide allows any user to use EASI-ORC, regardless of technical literacy. The guide includes technical instructions for the installation and application of each module.

To test EASI-ORC, we performed smFISH for the *SUC2* and *HSP150* mRNAs, which encode a secreted invertase protein necessary for sucrose utilization and a secreted *O*-mannosylated heat shock protein associated with the cell wall and its stability, respectively. We chose these mRNAs because they are known to localize to the ER^27^, making them good candidates for testing mRNA-organelle colocalization analysis. Our results show that EASI-ORC analysis was on par with manual analysis of the same data set, but with less variability since each run of the pipeline using the same parameters produces the same results. We show that EASI-ORC can perform its analysis using different smFISH probes and on different organelles (*e.g.* ER and mitochondria). It also can identify expected changes in localization stemming from differences in biological conditions. Importantly, it produces those results far faster than manual analysis can. In our testing it performed segmentation and spot identification for 100 different fields (each with 4 channels, 20 *z* planes, and up to 100 cells per field) in less than 24 hours. Once these steps are complete, mRNA-organelle localization analysis, including filtering, statistical analysis, and the creation of graphical outputs can be performed in minutes. Thus, the EASI-ORC pipeline provides a robust, approachable, and consistent solution for all the necessary steps in mRNA-organelle localization analysis. Each module can be used independently, with outputs allowing the user to perform troubleshooting at every step of the process. The entire pipeline’s code is open source, letting users change the system if they wish and our meticulously written step-by-step user guide allows any user, regardless of technical literacy, to use EASI-ORC for their data (scripts and user guide available on <u>the EASI-</u> <u>ORC GitHub</u>).

## Results

### Segmentation of Yeast Cell Images

The first stage of smFISH image analysis is cell segmentation, which outlines the perimeter of cells contained within the bright-field image (Figure 1A). Segmentation must be performed in a manner that enables the differentiation of yeast cells, one from the other, as well as from the background. To perform this, an ImageJ script was used to automate the YeastMate plugin^28^. YeastMate is a deep-learning based program that identifies yeast cells from bright-field or differential interference contrast (DIC) images. It is a fast and capable solution that works with a wide range of cell confluences (Figure S1).

The automation script produces binary black and white images corresponding to all images in a folder (the input folder). Producing an image that distinguishes between cells and background in a binary manner, will make the cell’s identification easier down the line. Since yeast cells in smFISH samples are fixed and stationary, their location in the *x* and *y* planes do not change across *z* planes. This allows us to select one *z* plane as the basis for the cell mask, thus, allowing for every cell to be identified across the different planes and imaging channels. YeastMate’s yield (*i.e.* percentage of individual intact cells identified within a field of cells) is dependent on the quality of the image - the correct focus and level of cell confluence. In an image with multiple planes, each plane will yield a different number of identified cells, even though they all essentially contain the same overall quantity. To maximize the data collected from each image, the *z* plane where YeastMate identifies the largest number of cells is chosen and duplicated to create the planes to create the final segmented image (*i.e.* the cell mask). Having the same mask in each plane allows one to keep track of each cell between the different planes. The product of this script is a folder with mask images that correspond to each raw image in the input folder.

### Organelle Segmentation

Organelle segmentation is a particularly difficult challenge when analyzing smFISH images, especially when the organelle in question is highly variable in shape across cells and experimental conditions (as with the ER, mitochondria, Golgi apparatus, lysosomes, and others). When performed manually, changing the image’s contrast and brightness between different cells may make the organelle appear larger or smaller than it actually is, due to the range of fluorescent signal intensities inherent to complex three-dimensional structures captured in two dimensions. The decision on the precise point in which an organelle starts and ends can be difficult to perform consistently and accurately. Even if an experienced researcher can perform the task well, marking the organelles in each cell and image becomes incredibly time-consuming. Simple computational tools, like those relying on signal strength thresholding (*i.e.* segmentation of an image into two parts according to a minimum and a maximum pixel intensity) often miss a large segment of the signals, due to the intrinsic signal intensity bias of thresholding. Put simply, thresholding ignores any signal beneath or above specific intensities, meaning any organelle represented by a range of intensities may not be well-segmented (*i.e.* incompletely delineated). Thus, automation at this stage of image analysis must be less biased towards general signal intensity and ideally be flexible enough to allow the user to segment different organelles. While deep learning is a popular means to approach similar tasks^31^, its use requires programming expertise not necessarily acquired by biologists. Also, these tools require large amounts of previously annotated datasets to train models for classification. Such fluorescent image datasets are laborious to produce and the ones available rarely fit the requirements of any given experiment. For example, a model trained to segment the ER won’t necessarily be reliable under different conditions (*e.g.* a mutation or growth conditions that affect ER structure). In contrast, simple image features (*e.g.* edge detection, texture, and intensity) can provide fast and robust segmentation by using classical machine learning tools that employ smaller datasets and less training for the segmentation of fluorescent images. Thus, we took advantage of the ImageJ plugin, Trainable WEKA Segmentation^29^,which comprises a set of tools utilizing machine learning and a simple-to-use graphical interface, allowing any user the ability to train a model for image segmentation that is outputted as a classifier file. Training can be performed quickly on a small scale (*i.e.* a few representative images usually suffice) to produce a classifier (model) for each organelle for each experiment. Once a classifier is trained, the EASI-ORC organelle segmentation script applies it for a folder of images (Figures 1B and 2A). The script produces a corresponding statistical map for each image in a folder. This statistical map gives each pixel in the image a normalized value between 0 and 1, corresponding to the probability that the pixel contains an organelle signal. This map can now be processed to a binary image using simple thresholding, *i.e.* using ImageJ’s default method, which is a variation of the IsoData^32^ method (Figure 2B). This two-stage approach of processing the raw image using a classifier, followed by thresholding the normalized statistical map, yields a mask that ignores noise and maintains the most usable signals.

**Figure 2.**
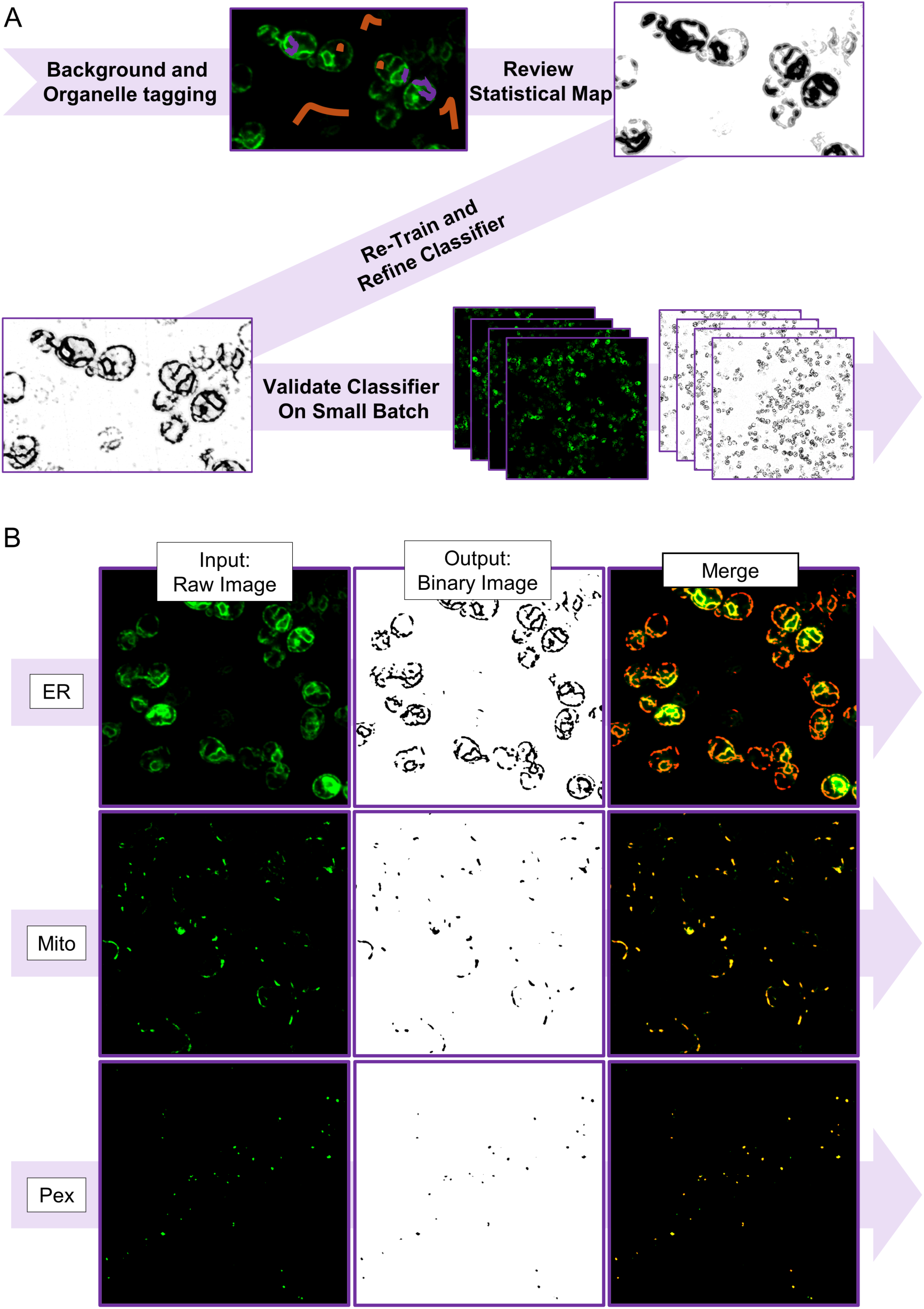
Organelle Segmentation. **A)** A diagram depicting the process of classifier training using the Trainable WEKA Segmentation plugin. The organelle (ER) is marked by a Sec63-GFP construct and image captured after smFISH (as described). The user tags the organelle (purple lines) and the background (orange lines) using the Trainable WEKA Segmentation GUI. A classifier is created based on the user’s tags. The output statistical maps designate a darker color per pixel, the higher the likelihood it contains an organelle signal (0 – white, 1 – Black). **B)** Left panels show examples of input images of the different organelles (shown in green), while and the middle panels display the binary image outputs produced by the organelle segmentation module of EASI-ORC (organelles are shown in black). The right panel shows the merge of the original organelle images and the binary mask images overlaid in red. Top row – ER marker (ER; Sec63-GFP); middle row - mitochondrial marker (Mito; Tom20-GFP); bottom row – peroxisomal marker (Pex; Pex3-GFP).

### Organelle Sub-segmentation Using a Proximity Marker

The delineation of an organelle often demands further sub-segmentation. For ER and RNA colocalization, for example, it is useful to divide the organelle into two distinct regions, due to differences in biological function. The perinuclear ER surrounding the nuclear membrane (nER) and the cortical ER, which extends from the nER to the plasma membrane and underlies the surface (cER)^33–36^. Defining the two sub-segments is another challenge when trying to automate the process of image analysis. In some cases, we may identify different segments of an organelle by their proximity to another marker (Figure 1D). In the case of the ER, proximity to the nucleus correlates with nER, while the lack of proximity correlates with cER locations. Therefore, a nuclear marker can be used as a proximity marker for the ER. By using the organelle segmentation module on a nuclear marker (*e.g.* DAPI or Hoechst), the ER can be easily sub-segmented. Any ER signal in proximity to the nuclear marker is designated as nER, while the remaining ER signal is designated as cER. The sub-segmented ER binary images are created in two steps. First, by subtracting the nuclear binary image from the ER binary image to produce a binary image of ER segments distant from the nucleus (*i.e.* the cER or distal mask) (Figure S2A). Second, the cER binary image is subtracted from the total ER binary image to produce the nER binary image (the proximal mask) (Figure S2B). This approach, exemplified by the ER and the nucleus, can be implemented using the same script for any sub-division of an organelle the user wishes to segment. This is provided that a suitable proximity marker which differentiates between the different sub-sections is available (Figure S2C). So, the EASI-ORC sub-segmentation module creates two binary images: one image corresponding to the organelle segments near a proximity marker and another image corresponding to organelle segments distal from the proximity marker. Similar to other modules, it also automates the process for a folder of images.

### smFISH Signal Identification

Many different computational solutions for the identification of smFISH signals exist. They allow for accurate and consistent signal marking, may offer a graphical interface to make the task more intuitive for users, and are often used to assist in the manual assessment of colocalization between two smFISH spots. The solution chosen for automation within EASI-ORC is RS-FISH^24^. It was chosen since it is ImageJ-based, relatively easy to automate, and due to its ability to use three-dimensional information in a multi-plane image to pinpoint a signal’s location at sub-pixel resolution. In addition, it allows the user to identify an appropriate signal threshold parameter in an interactive manner, making it useful across different imaging conditions. Thus, it provides an accurate and consistent designation of smFISH signal locations. The RS-FISH automation process requires the user to use the plugin manually once per signal source (*e.g.* RNA target or probe used) to ascertain the correct threshold (a simple process, explained thoroughly in the EASI-ORC user guide). Once the threshold is defined, the script can be applied to a folder of images. For each image, the script saves a table containing the *x-y-z* coordinates (width, height, and z-plane respectively), and intensity values for each smFISH signal identified (Figure 1C). These data are used in the EASI-ORC pipeline to identify the cellular localization that each smFISH spot belongs to and calculate the level of RNA-organelle colocalization (Figure 1E).

### Resolution of RNA-Organelle Colocalization

Performing RNA-organelle colocalization manually presents a very difficult challenge. The varied shape and intensities of organellar structures makes the decision-making process lean heavily on subjective parameters, which can lack consistency. Another facet of said challenge, is the fact it is time-consuming. Often, hundreds of cells are imaged for each sample and each of these cells may contain multiple FISH signals. Thanks to the segmentation methods used in EASI-ORC, the organelle borders and smFISH signals are well-defined (Figure 3A), and colocalization can be measured using simple tools available in ImageJ. As mentioned above, each cell is identified and delineated using the cell mask feature of YeastMate. The cells in the mask file are then enlarged slightly using the Dilate command to add pixels to their edges in order to improve coverage of peripheral organelles, such as cER. Any nearby cells are separated using the Watershed command in ImageJ. Next, the smFISH spot data is used to match the identified cells to their specific RNA signals. RNA-organelle colocalization is measured using a circle drawn with a user-selected diameter that is centered around the RNA spot’s location as identified by RS-FISH (Figure 3B). The circle is drawn on the organelle mask image(s) and the average signal intensity within the circle is measured. In case an organelle signal is found within the circle (*i.e.* within proximity to the smFISH signal), the average signal intensity will be greater than 0 and that spot will be designated as colocalized with the organelle. If the average intensity of the organelle signal is equivalent to 0, the spot is marked as not colocalized.

**Figure 3.**
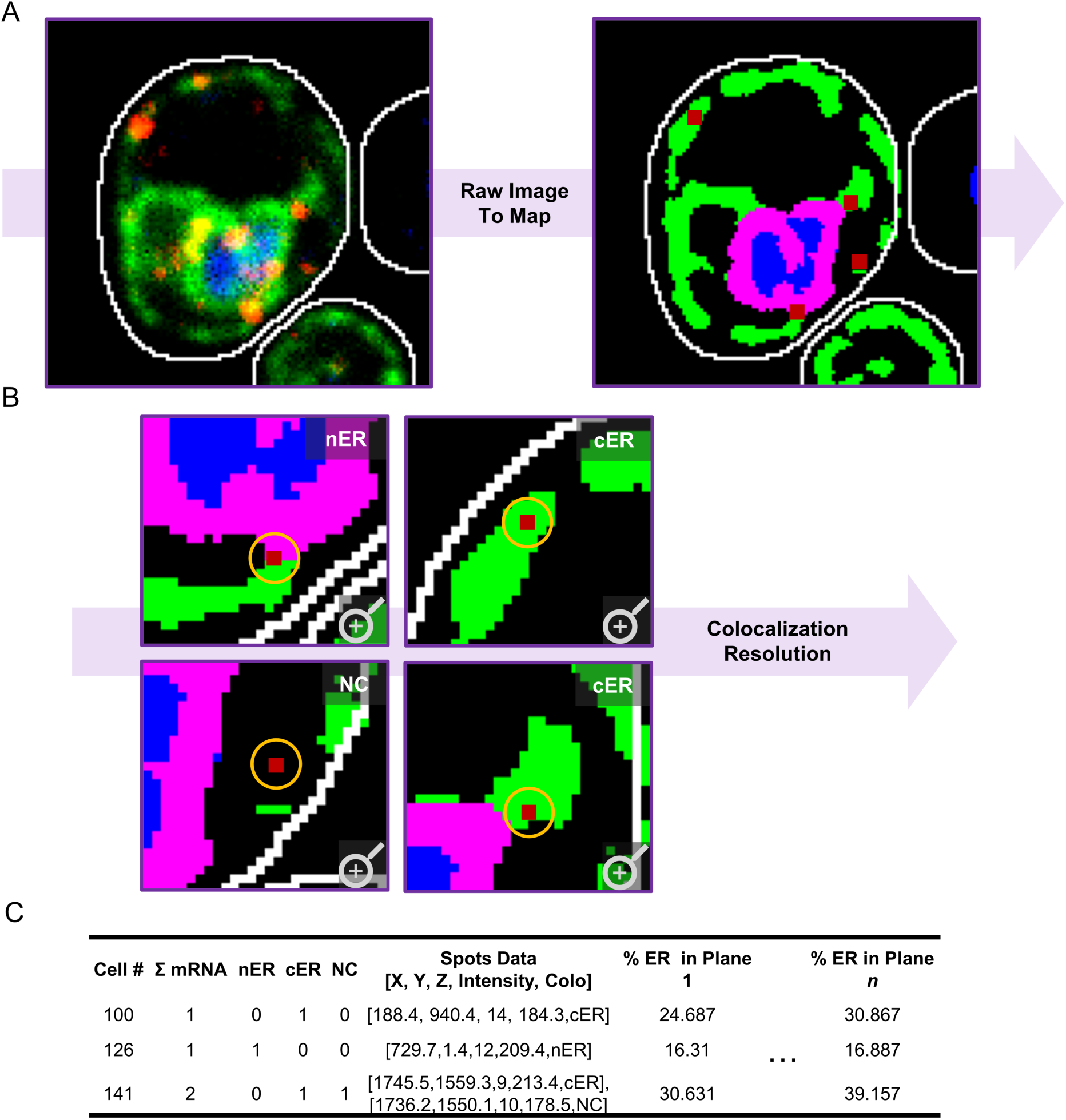
Resolution of Organelle-RNA Colocalization. After creation of the organelle and cellular binary maps, sub-segmentation (if needed), and smFISH spot identification, the EASI-ORC colocalization module resolves the colocalization status of each identified RNA spot. **A)** The lefthand image depicts an example of a raw image for a single cell. The ER is green (marked by Sec63-GFP), the nucleus is blue (dyed with DAPI) and the probes are red (*SUC2* probes labeled with TAMRA). The righthand image is a modeled representation of the cell, with different sub-cellular structures in different colors created from the produced masks. The cER is shown in green, the nER in magenta, and the nucleus in blue. The identified smFISH signals in the plane imaged (signals belonging to other *z*-planes not shown) are marked in red squares. White lines delineate the outline of the cell in both images. **B)** Colocalization resolution performed with EASI-ORC by measuring the organelle signal within a user-defined diameter. Zoom-in images of the binary map of a cell with red squares showing each identified smFISH signal are illustrated. The orange circle shows the area for organelle signal search for each smFISH signal, with the resolved pattern of colocalization written at the top right of each image (*e.g.* cER for cortical ER colocalization and NC for no colocalization). **C)** A representative table of the output as produced by the colocalization module. Each line contains data from a single cell, including the number of total mRNA spots, colocalized mRNA spots for each sub-segment of the organelle, and non-colocalized mRNA spots. A list containing the coordinates, intensity, and colocalization resolution of each cell is also created. In addition, each cell’s organellar coverage for every plane is measured and added to the table.

This module can work with an organelle that was sub-segmented into two parts. The user is prompted at the beginning of the run to choose between the sub-segmented and whole organelle mode. In case they choose the sub-segmented mode and to make sure that only cells where the sub-segmentation was successful are analyzed, the user must input a minimum value for the cellular coverage of the proximity marker in a plane (*i.e.* a fractional number between 0 for 0% and 1 for 100%). Any planes below the threshold will not be counted for colocalization calculations. Colocalization measurements are performed in the same manner as the whole organelle mode, with the addition of comparing the average signal intensity surrounding a spot for both sub-segments and selecting that of higher intensity. In the case of where the average intensities are equivalent, a new circle is drawn with a diameter of 1 pixel and the intensity is measured again. This is repeated with an increase of 0.5 pixels per iteration until the intensities of each organelle sub-segment are different, thus, resolving the specificity of colocalization accordingly.

The process is repeated for each spot identified and the data for each image is saved in a separate table (*e.g.* a csv file). Each row in the table represents a single cell and includes the data for the total number of mRNA signals identified in the cell and total of mRNA spots identified as either being colocalized or non-colocalized. Another column contains a list of coordinates, intensity, and assessment of RNA-organelle colocalization for each spot. In addition, a measurement of the relative size of the organelle signal (*i.e.* percentage of pixels that are designated as bearing the organellar marker in the mask) is also saved per each plane captured. These data can be used to remove specific planes (with their corresponding smFISH signals) that have an organelle signal too intense (*i.e.* “burned” signals that arise due to capture conditions or loss of cell integrity) or too small (*i.e.* lack of focus in a specific plane or loss of organelle marker).

### Data consolidation and analysis

Once image analysis using the EASI-ORC modules is complete, the raw colocalization data is available for study. We developed an online tool that allows analysis of the EASI-ORC output data using Python. The tool allows the user to upload sets of data, where each original image is represented by a single file and to name them as they wish. Once all data sets are uploaded, a consolidated table for each set is created and filtering can be performed. The toolset offers 3 filters (*e.g.* mRNA signal intensity, organelle coverage per *z* plane, and mRNA signals per cell) that need to be used successively. To demonstrate this, we employed data from an smFISH experiment using probes for *SUC2* mRNA and yeast strains expressing an ER marker fused to GFP (Sec63-GFP) (Figure 4). Each filter presents the user with a histogram output of the data that is updated dynamically, as the user changes the maximum or minimum values for it (Figure 4A). First, smFISH signals are filtered for intensity to allow removal of any outlier signals with intensity that is too high (*e.g.* a burned pixel) or too low (*e.g.* noise that was marked as a signal). Second, the *z* planes are filtered by organelle coverage (per cell) to remove specific planes. The relative organelle coverage in a cell changes according to organellar structure and experimental conditions, but in all cases extreme organelle coverage (*e.g.* due to over-exposure or cell inviability) will bias mRNA-organelle colocalization. A high value will yield a high colocalization fraction, while a significantly low value will yield the opposite. To avoid this bias, outlier *z* planes can be removed. Lastly, we provide a filter for the number of mRNA signals per cell. Similarly, outliers can be removed using this filter, but the values chosen should be based on the target mRNA’s expression, if known. For example, if the target mRNA is known to be of low abundance, it may be useful to ignore any cells that have an unusually great number of signals, since they may be an artifact and not representative of actual biology. For example, the results shown in Figure 4A, have been filtered (from top to bottom) for smFISH signal intensities from 150 to 600, for organelle (ER) coverages from 7% to 60%, and for mRNA signals per cell from 1 to 60.

**Figure 4.**
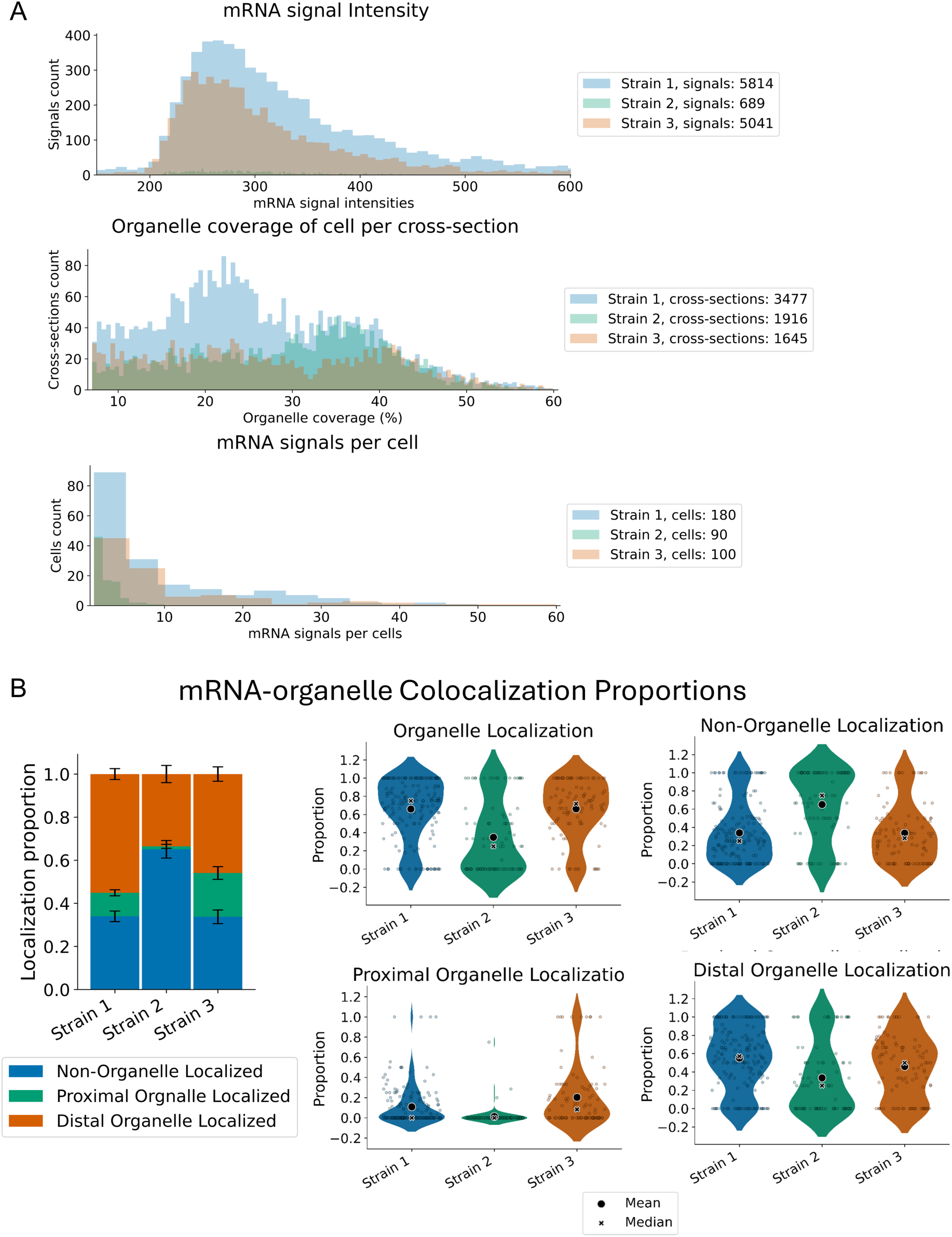
Data Filtering and Analysis. A toolset to analyze the EASI-ORC output data. Generic data labels are used as examples. Data produced from an smFISH experiment using strains with GFP labeled ER and SUC2 targeting probes marked by TAMRA. **A)** Dynamic filtering is performed using sliders to select minimum and maximum values for each filter. The data is presented graphically in histograms and updates as the user changes the filtering values. Filtering is performed successively by mRNA signal intensity, organelle coverage of a cell in specific planes, and the number of mRNA signals per cell. **B)** mRNA-organelle colocalization is presented using a stacked bar graph and violin plots. In the stacked bar graph, the proportion of each colocalization designation (non-colocalized or colocalized to each sub-segment of the organelle) can be seen in different colors. Errors bars represent the standard error of the mean for each. Violin plots for each colocalization designation include total organelle and non-colocalized data. When using the sub-segmented organelle mode, violin plots presenting the data for proximal and distal localized spots are also produced. Each dot in the plots represents a single cell. The black ‘X’ and circle represents the median and mean respectively.

Once filtering is performed, the graphical and statistical toolset can output a variety of plots to describe the data (Figure 4B and 5). Average mRNA-organelle colocalization proportions are calculated per cell and presented in a stacked bar graph, with standard error of the mean (SEM) error bars (Figure 4B). The proportion of mRNA-organelle colocalization can be presented in violin plots (Figure 4B). Each plot represents a single designation of colocalization (*e.g.* colocalized with the organelle, not colocalized, or colocalized with proximal or distal sub-segments) and each cell data point is represented by a dot (Figure 4B). The number of mRNA signals per cell (Figure 5A) and average organellar coverage per cell (Figure 5B) can also be presented in violin plots. Lastly (Figure 5C), scatter plots are used to present correlations between the proportion of colocalized smFISH-organelle signals (using total mRNA-organelle colocalization data) and either the average level of organelle coverage per cell or the number of mRNA signals per cell. The Pearson correlation coefficient is also calculated and presented in the plot (Figure 5C). Each figure can be edited by the user to change figure names, axis names, font sizes within the plot, and different plot styles. In addition, an image of the plot can be saved in several different file formats and quality presets, as chosen by the user.

**Figure 5.**
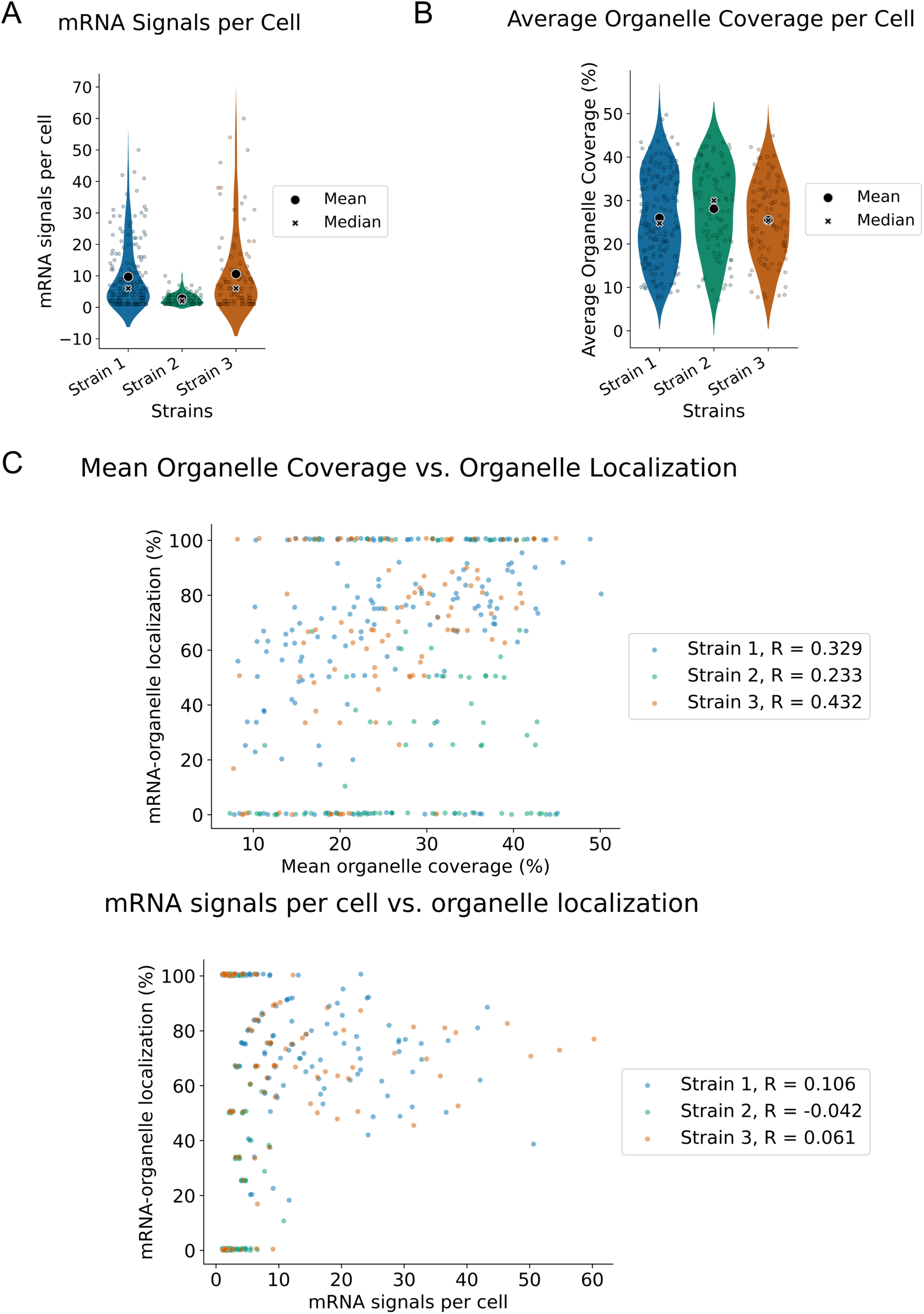
Other Analysis Results. **A)** The number of mRNA signals in a cell is presented using violin plots. Each dot represents a single cell. The black ‘X’ and circle represents the median and mean respectively. **B)** Average organelle coverage of the cell in the *z*-planes after filtering is calculated per cell and presented using violin plots. Each dot represents a single cell. The black ‘X’ and circle represents the median and mean respectively. **C)** Correlations between mRNA-organelle localization and either the mean organelle coverage or number of mRNA signals is presented in scatter plots. The Pearson correlation value is calculated and shown. Each spot represents a single cell.

Finally, a statistical summary table is produced. The table includes data on the number of cells identified, the number of cells remaining after filtering, Pearson correlation values and their statistical significance (p-value), mean colocalization proportions (with standard deviation and SEM), mean organelle coverage, and ANOVA test *p*-values between each data set (Table 1). All plots and tables created during the use of the analysis toolset can be saved and downloaded by the user.

**Table 1:**
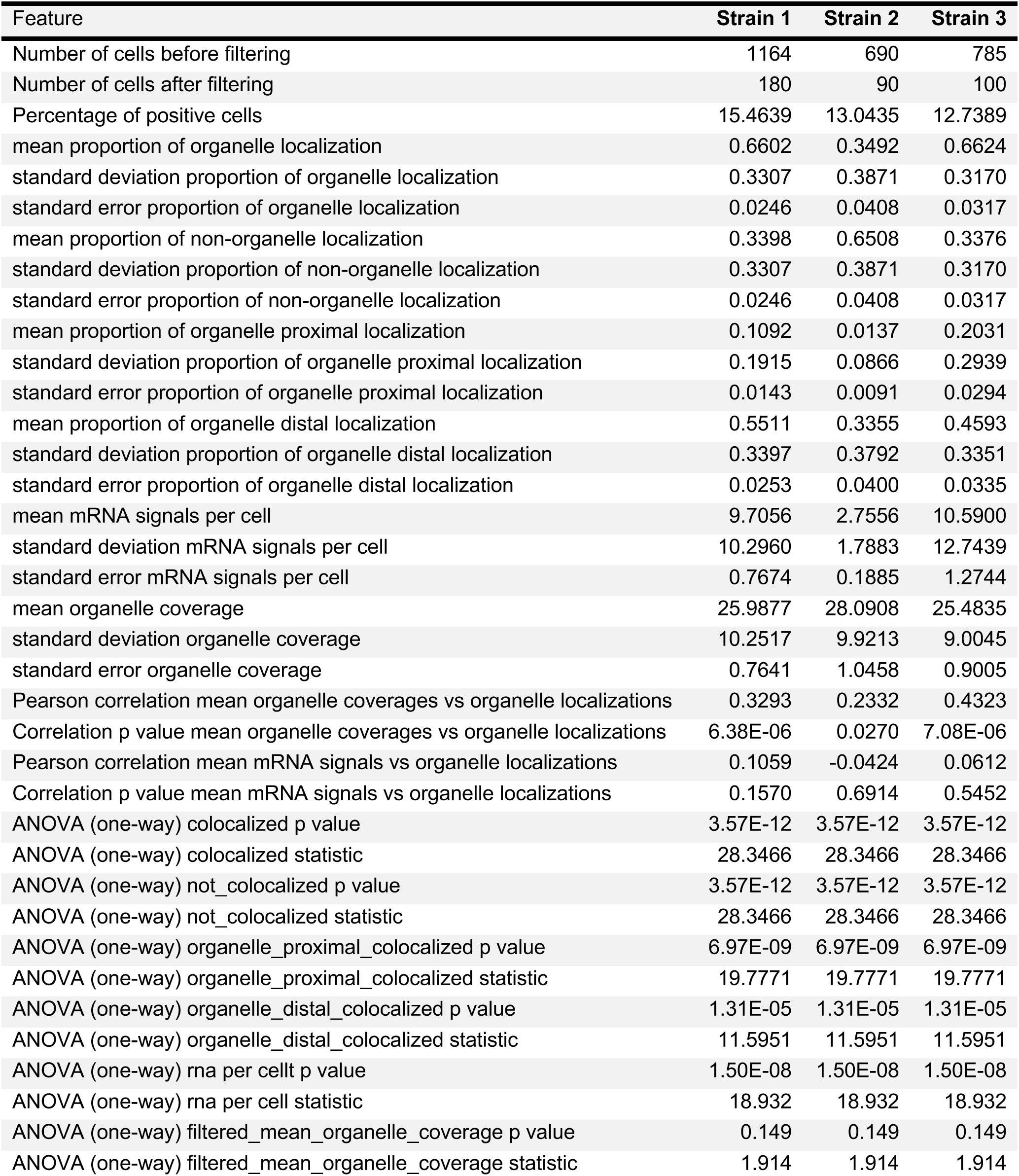
Example of Statistical Summary Table.

### Validation of the EASI-ORC analysis results

To confirm the functionality and accuracy of the EASI-ORC analysis pipeline, sample smFISH images were analyzed both manually and using EASI-ORC, and the mRNA-organelle colocalization results were compared (Figure 6A). For manual counting, RS-FISH was used for spot detection and colocalization with an organellar marker was judged by eye. In this experiment smFISH probes fluorescently labeled with Cy5 and complementary to *HSP150* mRNA were used. The organelles were marked by organellar proteins tagged with GFP at the carboxy-terminus: plasmid-expressed Sec63 for the ER and endogenously expressed Pex3 for peroxisomes (Pex3-GFP) or Tom20 for mitochondria (Tom20-GFP). *HSP150* encodes a secreted protein whose mRNA is ER-associated^27,37^. For 30 fields imaged, EASI-ORC analysis took approximately 8 hours, while manual analysis took at least twice as long. In addition, EASI-ORC analyzed >5-fold more cells than manual analysis (*i.e.* manual analysis was 44 cells per organellar marker, while EASI-ORC analyzed 262, 347, and 335 cells for the ER, mitochondria, and peroxisome, respectively) (Figure 6A). The results obtained between the procedures were highly similar. Manual counting showed that 46.99% of *HSP150* mRNA colocalizes with ER, 1.99% with mitochondria, and 0.87% with peroxisomes. Likewise, EASI-ORC analysis showed that 47.65% of *HSP150* mRNA colocalizes with ER, 4.21% with mitochondria, and 0.98% with the peroxisome (Figure 6A). Overall, *HSP150* mRNA-ER colocalization differed in proportion by only 1.6% (± standard error of 5.96%) between the two methods. For mitochondria, this difference was only 2.3% (± 1.02%) and 0.12% (± 0.6%) for the peroxisome.

**Figure 6.**
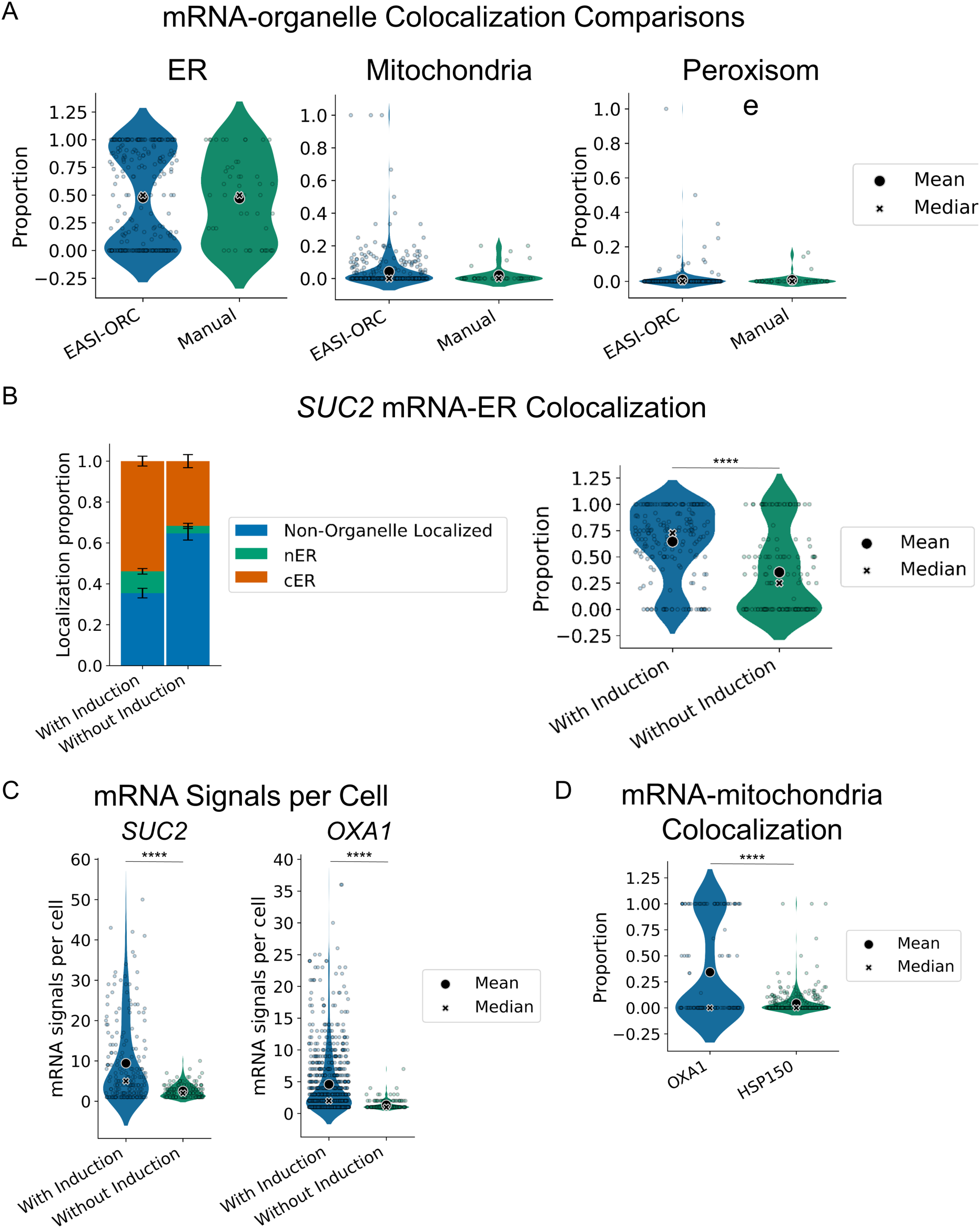
EASI-ORC validation. EASI-ORC results are highly similar to those obtained by manual counting. The figure shows the outputs from a series of experiments that measure RNA localization to the different organelles (ER, mitochondria, and peroxisome) using different probe sets, and identifies changes incurred due to varying biological conditions. **A)** EASI-ORC outputs the results of mRNA-organelle colocalization analysis for the ER, mitochondria and peroxisomes (based on one biological repeat) using Cy5-labeled probes targeting *HSP150* mRNA. Each sample was also analyzed manually, using RS-FISH for spot identification (number of cells scored: manual analysis n=44 per experiment; EASI-ORC analysis, ER n=262; mitochondria n=347; peroxisome n=335). Any differences between EASI-ORC and manual analysis were not significant according to the ANOVA test (not shown). **B)** EASI-ORC outputs results corresponding with different treatments, as shown using *SUC2* induction under low glucose growth conditions (*e.g.* 0.05% for induction and 2% for no induction for 2 hrs; one biological repeat). Bar graphs showing mRNA-ER localization percentages with sub-segmentation to nER and cER. Total mRNA-ER localization presented in violin plots (ANOVA test *p*-value = 8.49×10^−13^; number of cells scored with induction n=204; without induction n=142). **C)** The number of *SUC2* mRNA signals per cell shows a significant decrease (ANOVA test *p*-value = 4.11×10^−14^) without induction. A separate induction experiment was performed for *OXA1* (cells grown for 1.5 hours in 3% glycerol- or glucose-containing medium) also shows a significant decrease on mRNA levels (ANOVA test *p-*value = 2.31×10^−14^; number of cells: with induction n=838, without induction n=148). **D)** mRNA-mitochondria localization of both *OXA1* and *HSP150* mRNA was compared. *OXA1* mRNA showed a significantly higher colocalization levels (ANOVA test *p*-value = 2.98×10^−27^; number of cells: *OXA1* mRNA, n=148; *HSP150* mRNA, n=347).

To demonstrate EASI-ORC’s ability to identify changes in mRNA-organelle localization under different conditions (Figure 6B), a comparison was performed between yeast cells grown on low glucose to induce the *SUC2* gene (*e.g.* 2hrs growth in 0.05% glucose) to those grown under *SUC2* repressive conditions (*e.g.* 2% glucose)^38,39^. The level of *SUC2* mRNA-ER localization (for both nER and cER) decreased from 64.6% on low glucose to 35.3% on high glucose. Changes in the number of mRNA signals per cell upon induction were also measured for *SUC2* (Figure 6C). A significant increase in the number of mRNA signals per cell was found, with an approximately 4-fold increase (*e.g*. 9.41 signals per cell with induction and 2.57 signals without) (Figure 6C). An induction experiment was also performed for *OXA1* mRNA (*e.g.* 1.5 hrs growth in 3% glycerol instead of glucose). Again, with no induction, a significant decrease in mRNA signals was observed (*e.g.* 4.59 signals per cell with induction and 1.35 signals without, Figure 6C). Lastly, we tested EASI-ORC’s ability to compare mRNA-mitochondria localization between two different mRNAs: *OXA1* and *HSP150* (Figure 6D). In both cases, Tom20 was labeled fluorescently with GFP and smFISH probes for each target were labeled with Cy5. *OXA1* mRNA localization to the mitochondria was significantly greater than that of *HSP150* (*e.g.* 34.28% and 4.25% respectively).

## Discussion

Our work demonstrates that EASI-ORC is a useful set of analysis tools that provides a reliable, consistent, and efficient solution for smFISH colocalization analysis in yeast. The expansive user guide we provide goes into meticulous detail (available on <u>the</u> <u>EASI-ORC GitHub</u>) to help bypass technical challenges that researchers may face. It explains, step-by-step, how to use each of EASI-ORC’s modules, *i.e.* the preparations required before using each module and considerations that must be made when choosing important input and filtering values. It also includes a thorough explanation of all steps taken by each script. The same user guidance applies to EASI-ORC’s statistical analysis and graphing tools, with clear instructions presented for every step.

EASI-ORC performs the cellular segmentation of yeast and successive sub-cellular segmentation of any structure (*e.g.* organelle) to measure mRNA-organelle colocalization. Theoretically, it can be utilized for other types of cells by replacing its cell segmentation step with an appropriate solution that outputs binary images of any organism. Also, thanks to the use of a dynamic user-trained organelle segmentation method, it is capable of performing colocalization with any organelle or other sub-cellular structure and can even sub-segment an organelle to two different sections (providing a proximity signal exists). Our testing shows it produces results highly similar to those obtained using manual analysis, however, using significantly less time and effort, yet while taking into account significantly more cells (Figure 5A). It was also shown to accurately and effectively measure biological changes and differences in mRNA-organelle localization and mRNA abundance. Induction of *SUC2* led to a significant increase in mRNA-ER localization proportions (Figure 5B), that is consistent with prior findings^38,39^. In addition, an increase was observed in *SUC2* and *OXA1* mRNA abundance upon inducive conditions (Figure 6C). EASI-ORC analysis showed that a ∼4-fold increase in *SUC2* mRNA abundance is induced under low glucose conditions, similar to that shown by others^40^. *OXA1* mRNA levels were increased ∼3.5-fold upon growth in glycerol-containing media, as expected from the increase in mitochondrial activity under respiration conditions (Figure 6C). Also, when comparing *HSP150* mRNA-organelle colocalization, a greater level of ER colocalization was registered, when compared to either the mitochondria or peroxisome (Figure 6A). This was expected due to Hsp150 being a secreted protein^41^ and *HSP150* mRNA showed similar ER localization patterns in previous studies^27^. In addition, mRNA-mitochondria localization was compared between *HSP150* and *OXA1* mRNAs. *OXA1* mRNA showed a ∼6-fold greater proportion of localization to the mitochondria than *HSP150* mRNA (Figure 6D). The latter preferentially localizes to the ER, while *OXA1* mRNA is known to localize to mitochondria^42^. To alleviate any concern that organelle coverage correlates with the level of mRNA-organelle colocalization, we examined colocalization per cell with the average organelle coverage per cell (across *z* planes). We found no correlation between organelle size and *HSP150* mRNA-organelle colocalization levels in any of our experiments. Pearson correlation values between average organelle coverage (for all *z* planes remaining after filtering) were −0.062, 0.023 and −0.077 for ER, mitochondria and peroxisomes respectively (Figure S4). Thus, the increase in colocalization levels is not a product of organelle size.

In addition to the script codes and user guide, we provide the models used by us to segment ER (using the Sec63-GFP marker), nucleus (using DAPI) peroxisomes (Pex3-GFP) and mitochondria (Tom20-GFP), in hopes they will be useful as preliminary models for other EASI-ORC users. Also, our pipeline automates the process of identifying smFISH signals from multiple images. Notably, each EASI-ORC module can work independently of any other, allowing the user to pick and choose the modules useful for them. This also allows implementation of changes and upgrades relatively easily.

In conclusion, EASI-ORC performs colocalization assessment significantly faster and more consistently than the arduous process of manual analysis. In the future, we hope to continue development of the EASI-ORC pipeline and add features such as automation for additional types of cell segmentation or two-spot colocalization, as well as adding more filtering options for the statistical analysis module. Beyond image analysis, EASI-ORC provides fundamental statistical analysis of mRNA-organelle colocalization, with dynamic graphical filters and easy-to-edit plotting tools. In its current form, EASI-ORC is a free and accessible automated solution for the analysis of common FISH assays that are usually performed manually. The pipeline allows the user to go from raw images to data visualization relatively quickly and the user guide makes EASI-ORC profoundly easy to use. We believe it facilitates the removal of technical and logistical barriers for RNA-organelle colocalization assays and improves both the accuracy and consistency of small- and large-scale FISH experiments.

## Methods

### EASI-ORC pipeline modules

The pipeline is divided into 6 modules (Figure 1) for yeast cell segmentation (Module 1), organelle segmentation (Module 2, organelle sub-segmentation (Module 3), smFISH signal identification (Module 4), colocalization resolution (Module 5), and data consolidation and analysis (Module 6).

The yeast cell segmentation module (Module 1) uses brightfield or DIC images of yeast to segment cells from the background. It automates the ImageJ plugin YeastMate^28^ for use on a folder of images and outputs a mask image for each image provided.

The organelle segmentation module (Module 2) uses images of fluorescently labeled sub-cellular structures (*e.g.* arising from fluorescent protein-tagged organellar marker proteins) to segment them from the rest of the image. By automating the Trainable WEKA Segmentation ImageJ plugin^29^ and using a user-trained model, it performs segmentation on all images in a folder and outputs statistical map and mask images for every image provided.

The organelle sub-segmentation module (Module 3) uses a proximity marker to sub-segment an organelle into two parts. This is useful in cases different parts of an organelle have different biological roles (*e.g.* the ER has two distinct sections called the perinuclear and cortical ER). The proximity marker is any imaged structure that designates different parts of an organelle on the basis of proximity to one distinct organellar sub-section. Any organellar signal overlapping the proximity marker is defined as a proximal section and any sections that do not are defined as distal sections. By using image masks of both the organelle and proximity marker, and employing simple subtraction, separate masks of the two sub-segments (*i.e.* proximal and distant) are produced. The proximity marker mask is subtracted from the organelle mask to create the distal mask image and the distal mask is subtracted from the organelle mask to produce the proximal mask image.

The smFISH signal identification module (Module 4) automates the use of RSFISH^24^ to identify smFISH probe signals in each image within a folder. It requires the user to identify the RSFISH threshold parameter and outputs a table containing all the identified RNA spots in an image. Each row represents a single spot and contains the *x*, *y*, *z* coordinates (width, height and z-plane, respectfully) and the spot’s intensity.

The colocalization resolution module (Module 5) uses the mask images and spot data tables produced by the other modules to associate each smFISH spot with a specific cell and resolve its designation of being co-localized (or not) with an organelle. The module can work with either a whole organelle or a sub-segmented one. A colocalization designation includes colocalization with the proximal, distal or whole organelle (*i.e.* in case no sub-segmentation was performed), or not colocalized at all. The resolution of RNA-organelle colocalization is performed by examining a surrounding circle around each smFISH spot, as determined by RSFISH, and using a diameter provided by the user (*e.g.* a diameter of 1, meaning a signal less than one pixel away; see User Guide). The signal diameter should be decided by the user, after closely observing several signals in their images. The size of an smFISH signal depends on the probe set used, the target mRNA and image capture conditions. If an organellar signal is present within the circle, the spot is resolved as being colocalized. If there is no signal present, it is deemed not to be colocalized. The module outputs a table for each image provided, with the data derived from each cell and represented by a single row. This includes data regarding the number of mRNA spots per cell, the number of mRNAs with each colocalization designation, the coordinates and intensity for each RNA spot, and the level of organelle coverage within the cell per *z* plane.

The data consolidation and analysis module (Module 6) is a toolset written in Python that takes the tables produced by the colocalization resolution module and consolidates them according to treatment or strain (defined according to the order the user uploads the table files). It allows for filtering by signal intensity, organelle coverage, and the number of smFISH signals per cell. After filtering the user may produce graphs and statistical analysis data, or perform the analysis by using the filtered output tables.

### Yeast strains and growth conditions

Parental yeast strain for all studies is BY4741 (*MAT*a*, his3Δ1, leu2Δ0, met15Δ0, ura3Δ0*). Cells were grown in liquid or on plates of YPD (1% Yeast extract, 2% Peptone, 2% Dextrose) at 30℃, unless mentioned otherwise.

### Plasmids used in this study

Sec63-GFP: Plasmid containing Sec63-GFP under the control of the T7 promotor and bearing a *URA3* selection marker.

### smFISH Probes used in this study

FISH probe sets targeting *SUC2*, *HSP150* (Biosearch Technologies, Novato, CA), and MS2V3^30^ (Sigma-Aldrich, St. Louis, MO).

### smFISH procedure

Yeast expressing organellar markers (*e.g.* Sec63-GFP from a plasmid, and Tom20-GFP or Pex3-GFP tagged endogenously) were grown to mid-log phase in YPD. When *SUC2* was the target mRNA yeast were subjected to *SUC2* induction of 1.5 hours in low glucose synthetic media containing 0.05% glucose (*e.g.*, synthetic complete). Crosslinking was performed in the same medium upon the addition of paraformaldehyde (4% final concentration), and incubated in room temperature for 45 minutes, with rotation. Cells were washed 3 times with ice-cold Buffer B (0.1 M potassium phosphate buffer, pH 7.5, containing 1.2 M sorbitol). Cells were spheroplasted in 1 ml spheroplast buffer (Buffer B with an addition of 20mM ribonucleoside vanadyl complexes (VRC) (Sigma-Aldrich), 20 mM β-mercaptoethanol, and lyticase (Sigma-Aldrich) (5 U per O.D.600 unit of cells)) for 10 min at 30°C. Spheroplasts were centrifuged for 5 minutes at low speed (3500 rpm) at 4°C and washed twice with ice-cold Buffer B. Washed spheroplasts were resuspended in 1 ml cold 70% ethanol and incubated overnight in −20°C. After which, 250 µl of each sample was pelleted and washed twice with 1 ml 2x SSC buffer (0.3 M sodium chloride, 30 mM sodium citrate) at room temperature. Samples were then washed once with 10% formamide in 2x SSC buffer with 15 min incubation at room temperature. Samples were pelleted and 1-2 ng of fluorescent oligonucleotide probe was added in 50 µl of buffer H (10% formamide, 10 mg/ml *E. coli* tRNA, 2x SSC, 2mg/ml bovine serum albumin, 2 mg/ml VRC, 10% dextran sulfate) and incubated for 2 hrs at 37°C in the dark. Following that, 500 µl of 37°C 10% formamide in 2x SSC buffer was added to each sample, followed by gentle mixing, before incubation for 5 min in 37°C in the dark. Samples were then pelleted and washed with 10% formamide in 2x SSC with incubation of 5 minutes at 37°C in the dark. Samples were pelleted once again and resuspended in 0.1% Triton X-100 in 2x SSC and incubated at room temperature for 5 min in the dark. Samples were pelleted and resuspended in 500 µl DAPI (0.5 µg/ml) in 2x SSC and incubated for 2 min at room temperature in the dark. Samples were pelleted and washed with 2x SSC. Lastly, samples were resuspended in 100-200 µl 2x SSC and 5-10 µl were put on microscope slides pre-prepared with 0.01% poly-L-lysine (Sigma-Aldrich). A cover was placed and samples were taken to imaging on the same day. Samples were imaged using PCO-Edge sCMOS camera controlled by VisView installed on a VisiScope Confocal Cell Explorer system (Yokogawa spinning disk scanning unit); CSU-W1 and an inverted Olympus microscope (×100 oil objective; excitation wavelengths: GFP – 488 nm; TAMRA – 555 nm; Cy5 – 640 nm). 12-25 z-planes were captured, with 0.2-0.3 z-step sizes. Image analysis was performed with EASI-ORC, as described.

## Supporting information

Easi-Orc User Guide

## Supplementary Figures

**Figure S1.**
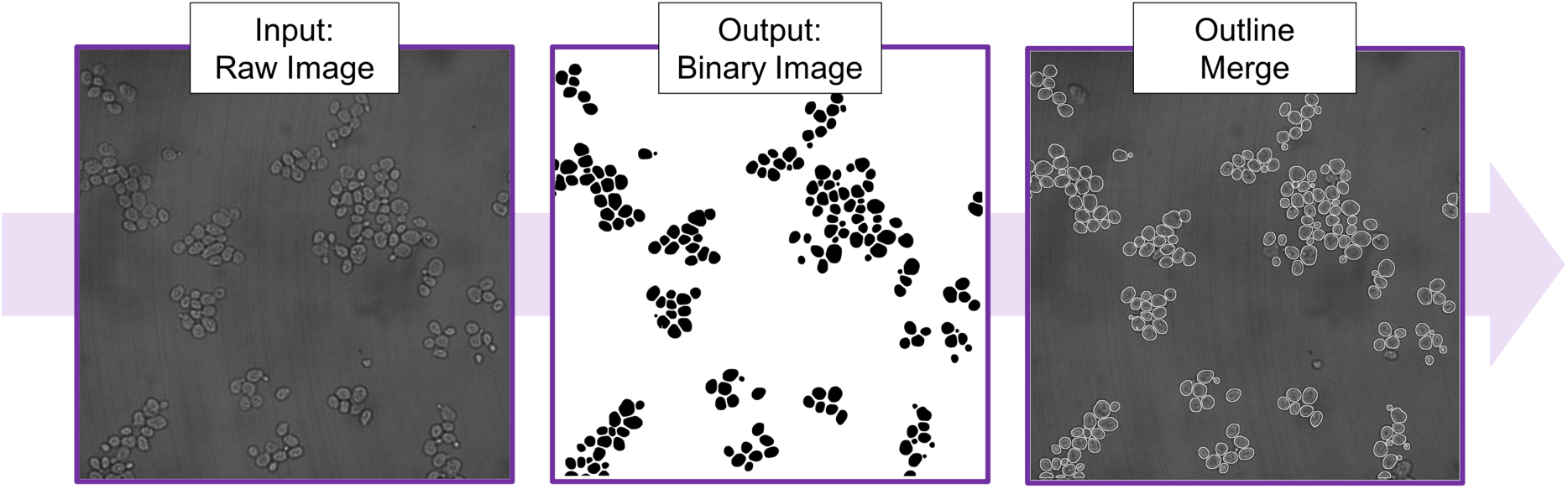
Yeast Cell Segmentation with YeastMate. Using YeastMate, the cell segmentation module identifies the cells in raw brightfield or DIC images and outputs a binary image of the cells. In the merged image, the binary image is used to outline each identified yeast cell.

**Figure S2.**
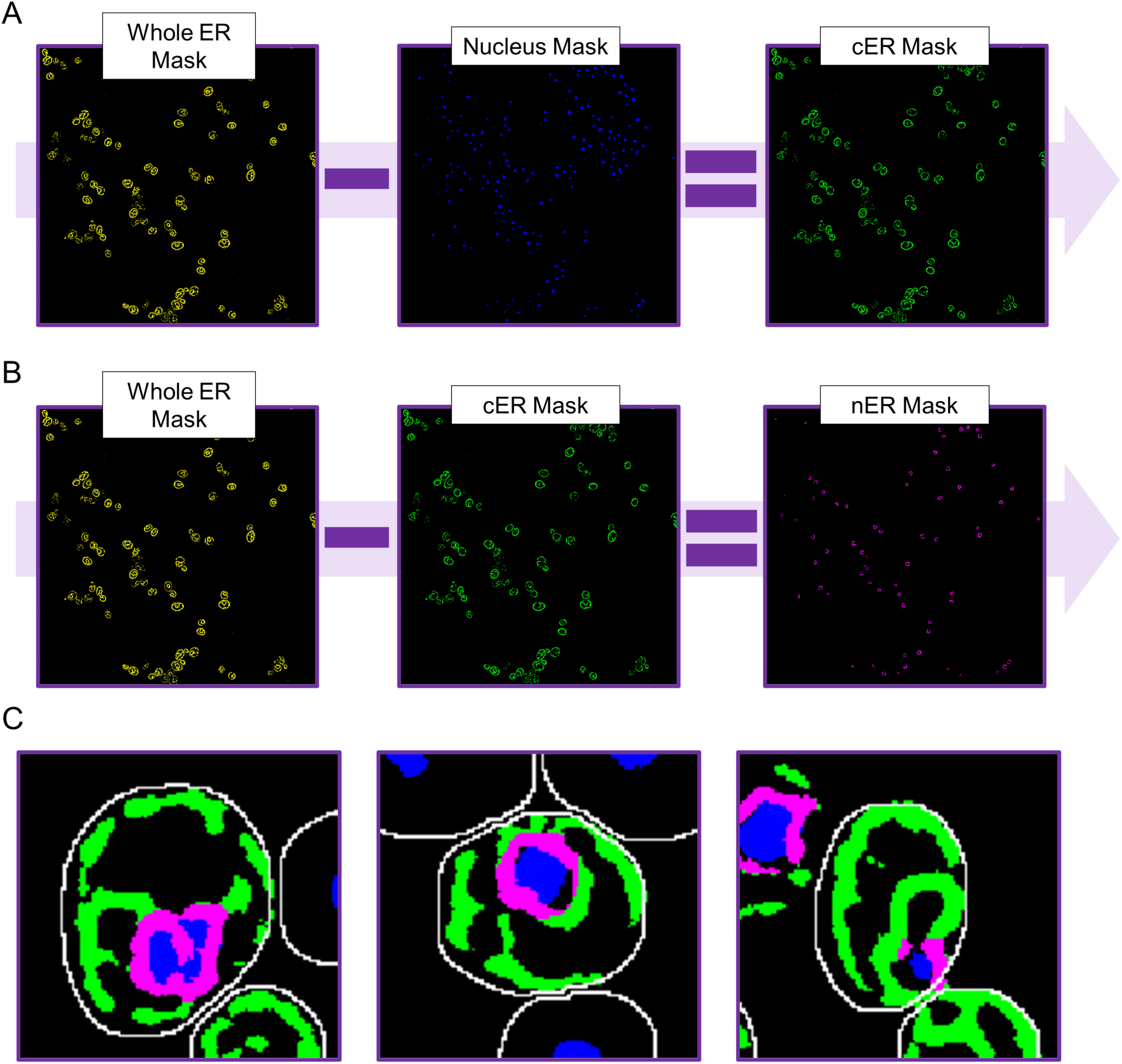
Organelle Sub-division. Organelle sub-division exemplified using ER and nuclear masks produced using the organelle segmentation module of EASI-ORC. A) A nuclear mask (in blue) is subtracted from a whole ER mask (in yellow) to produce a cortical ER (cER) mask (in green). The nucleus mask goes through three iterations of dilation (i.e. addition of pixels to the edges) to improve its coverage of the nuclear ER (nER). B) The produced cER mask is subtracted from the whole ER mask to produce the nER mask (in magenta). C) Organelle sub-segmentation is dependent upon proximity marker accuracy. From left to right are examples for good, partial, and insufficient proximity marker coverage. cER in green, nER in magenta, nucleus in blue. The white outline delineates the cell.

**Figure S3.**
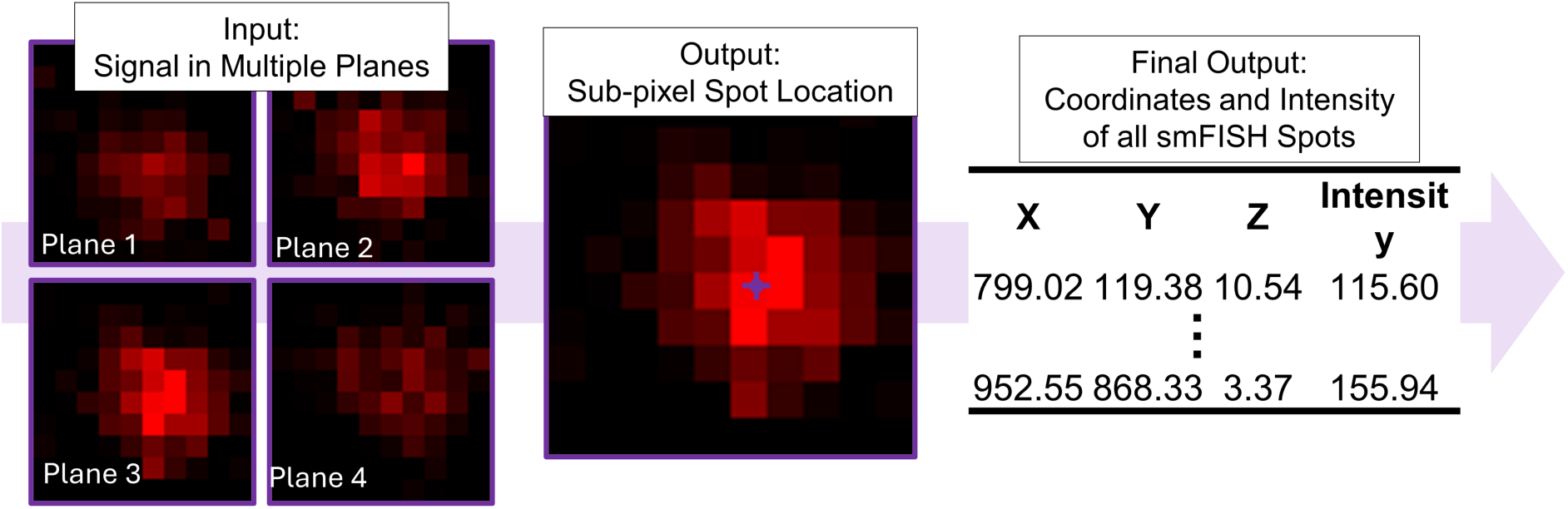
smFISH Spot Identification. RS-FISH uses data from all planes available in the image to identify the sub-pixel location of each spot. An output table is produced for each image, containing location and intensity data of all identified spots. On the left, a zoomed-in image of a spot in multiple planes (as detected by SUC2 probes labeled with TAMRA). In the middle, the plane with the most intense spot is presented with the sub-pixel location identified by RS-FISH. On the right, an example table of the EASI-ORC spot identification module output produced for each image. The table contains coordinates (X for width, Y for height and Z for plane) and intensity of all spots identified in the image.

**Figure S4:**
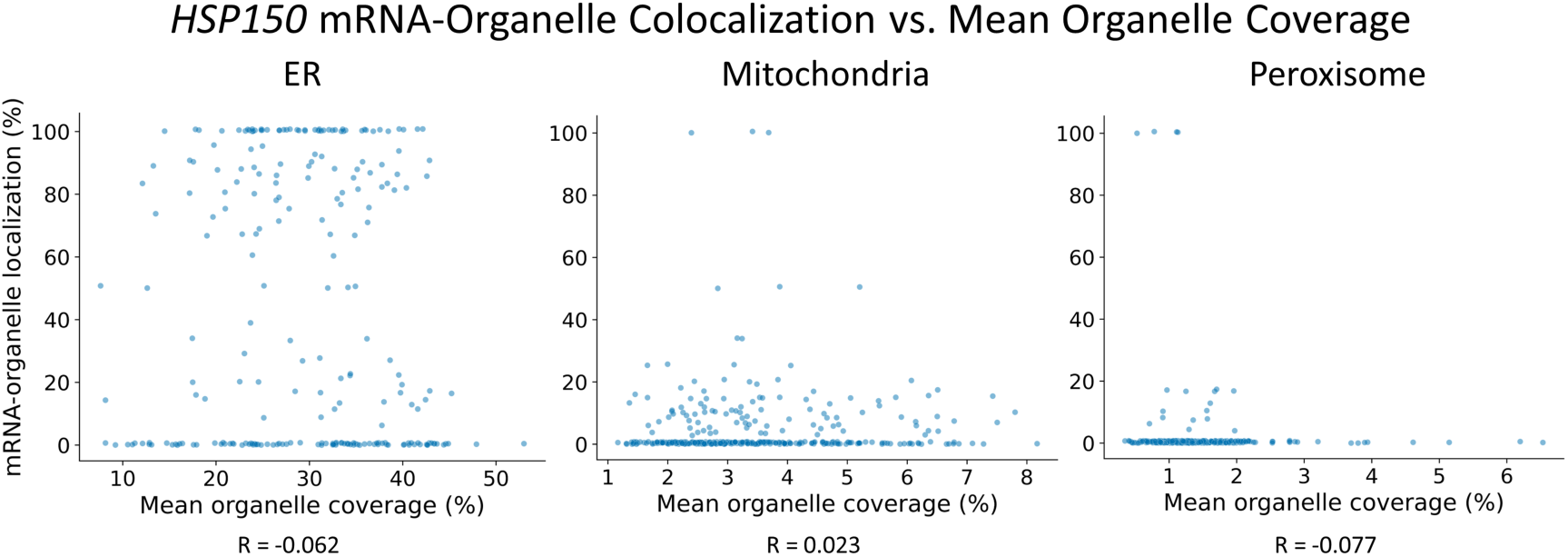
*HSP150* mRNA-Organelle Colocalization vs. Mean Organelle Coverage. Scatter plots representing the relationship between *HSP150* mRNA-organelle colocalization and mean cellular organelle coverage. Each spot represents a single cell. Data for the ER, mitochondria and peroxisomes (based on one biological repeat) was collected from smFISH experiments using Cy5-labeled probes targeting *HSP150* mRNA (number of cells scored: ER n=243; mitochondria n=347; peroxisome n=328).

