## Supplementary material for "EASI-ORC: A Pipeline for the Efficient Analysis and Segmentation of smFISH Images for Organelle-RNA Colocalization Measurements in Yeast": Easi-Orc User Guide

### EASI-ORC Guide:

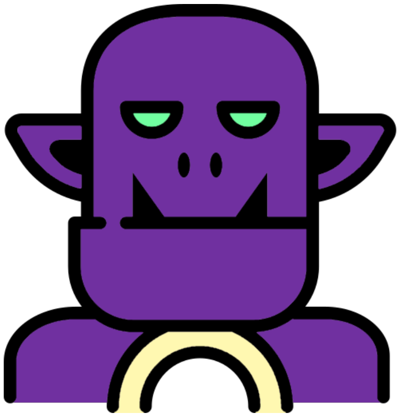

#### Introduction

This document explains the different steps required for fluorescent image analysis using EASI-ORC and makes them accessible with relatively little prior know-how. It requires the basic knowledge of working with computers and how to activate scripts in Fiji (Plugins → Macros → Run…). Prerequisites, such as file formats or needed values are explained in each section, so please read the instructions carefully. Examples shown in this guide were mainly based on ER segmentation, but the processes described can also be used for other organelles. The guide and scripts were written and employed using a Windows-based PC (Windows 11). Use of any of the scripts on different systems may create issues. All script codes can be found in the EASI-ORC GitHub. Each module describes the steps taken by the corresponding script. These descriptions can be used to manually perform the steps through FIJI's menus, if you wish to check the effect each step has on your images.

**Applications you need to install before beginning:**

Fiji can be downloaded for free here.

YeastMate standalone app can be downloaded for free here (download .exe file)

YeastMate plugin Fiji installation can be performed by following instructions on this page.

RS-FISH plugin Fiji installation can be performed by following instructions on this page.

#### 1. Requirements and Folder Structure

By and large, EASI-ORC can work on any modern Windows-based system. The faster the CPU the faster it will work, but it can complete its work without needing state-of-the art hardware. However, it is limited by RAM size. The higher the image resolution, the more RAM is required. The system was tested using 2048x2048 images and for that resolution we would recommend at least 32 GB of RAM. Anything less can result in some of the steps (*i.e.* cell and organelle segmentation) failing to reach completion.

Another requirement concerns the way the images are captured. smFISH signals are identified using data from multiple fields and planes of capture. As such, making sure that your fields are correctly focused is important in order to maximize the number of usable cells.

Specifically, signals should be least focused on the first and last *z* planes and most within the middle of the range.

EASI-ORC uses a specific folder structure. Every image from each channel captured must be saved in a separate folder that contains only images. The images should be in the TIF format. In the next segment we include a script that can convert the images to TIF format and order them in folders according to the channel. Lastly, high resolution images require a large amount of space. Before running EASI-ORC, make sure you have available space that is at least 3 times that of the total of your raw images. After completion of the scripts and image analysis, you can remove any images you wish, unless you plan to repeat the colocalization analysis.

The folder structure created by EASI-ORC is presented in the figure below.

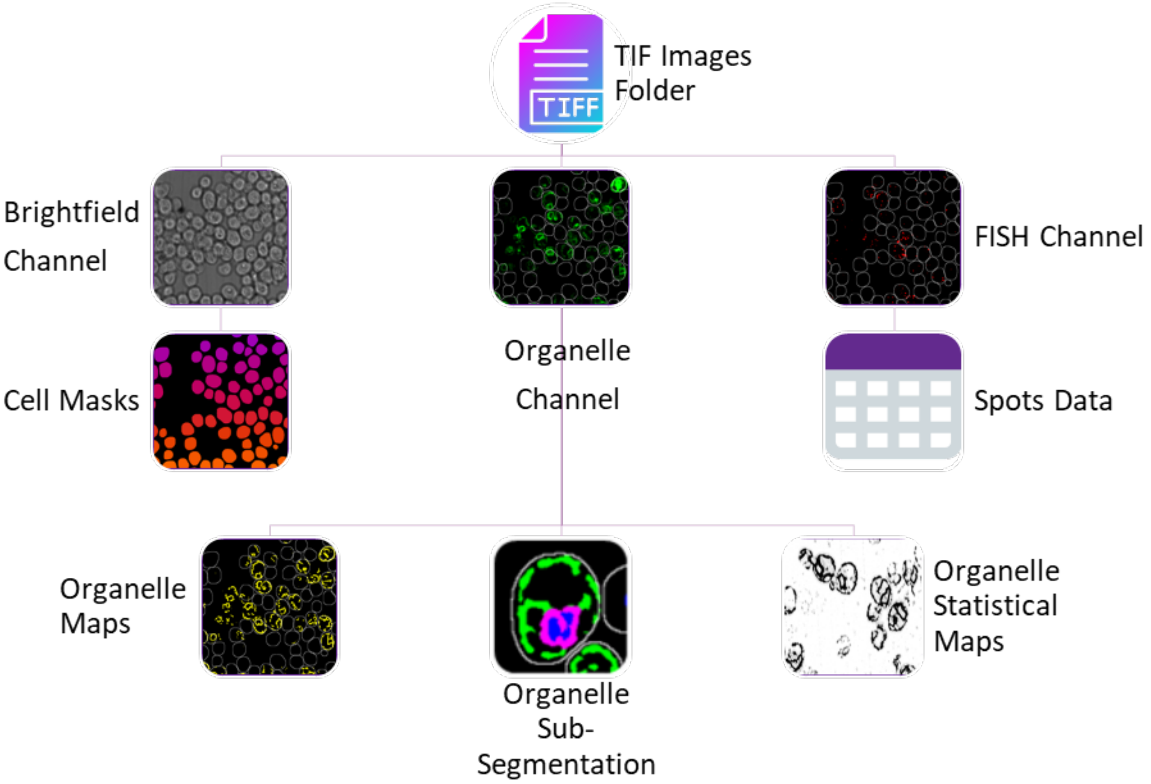

#### 2. How to use a Fiji Script

Each script is written as a single 'ijm' file. To use a script, simply open Fiji and drag the file icon to the Fiji menu window. A new window will open that contains the script's code. Press the 'Start' button (located below the code section of the window) and the script will run. Any inputs needed will appear in pop-up windows.

#### 3. Convert Raw Images to TIF and Arrange in Folders

Most microscope software outputs the images in a format unique to the manufacturer. These file types (*i.e.* bio-formats) can be less accessible for many programs or processes. Accordingly, we must make sure all files are in the correct format. This script (tested mainly on 'stk' files should also work with other bio-formats) automates this process. This script assumes each channel and field are saved in a separate image file (*i.e.* no multi-channel images). File names must be unique for each channel. It can have the name of the marker (*e.g.* GFP, DAPI, mCherry, etc.) or an arbitrary name (*e.g.* c1, c2, etc.). The user must know these names and input them (case-sensitive) once they run the script.

**Important** – if your files are formatted differently (in a multi-channel image, for example) this script won't work. You must first convert each channel to a separate image. You must have each channel's images collectively placed in separate folders, in TIF format, before using the EASI-ORC scripts.

**Script Steps:**

1) The user will be asked to input the directory containing the images and their file type (this usually appears at the end of the file name, after a dot).
2) The user will be asked to input the number of channels, as well as each channel's name. Channel names must be the same as they appear in the corresponding file's name (**case-sensitive**) and they must be unique per channel.
3) Each raw image will be opened sequentially and saved in its proper folder as a 'tiff' file, with its original name.
4) The 'tiff' images will be saved in a new folder located in the same location as the raw images. The main folder is called 'Tiff Images' and sub-folders for each of the channels will be created in it.

#### 4. Use YeastMate to Segment Yeast Cells in an Image

One of the first challenges in image analysis is identifying and defining each cell in the image. When it comes to images of yeast cells, YeastMate is a Fiji plugin that provides fast and trustworthy results using a pre-trained, machine learning approach. The script uses the app's default settings, with a threshold value of 0.6 to maximize the number of identified cells.

To use the YeastMate plugin, you must first install its standalone application. Failing that, the script described here will not work. The plugin itself must also be installed in Fiji. To choose the best *z* plane from a multiplane image, masks of each plane will be created and the one containing the largest number of cells will be chosen for the final image. In addition, YeastMate must be run on a computer without interruption. It can take a long time to get a mask when using high resolution images with many *z* planes. Thus, if you have many images to process, please take that into consideration. For example, when processing 50 images with 12 *z* sections each at 2048x2048 pixel resolution, the script needed approximately 4 hours to complete. Timing will vary greatly according to available computing resources and image size.

Once you install the YeastMate application and Fiji plugin, you can start the application to run the EASI-ORC module.

Follow these steps:

1) The program will start in 'Start a new job' tab (this and the following stage can be performed before you run the script).

2) Press the ‘Detection’ button under the settings. Your settings should be all be turned off,

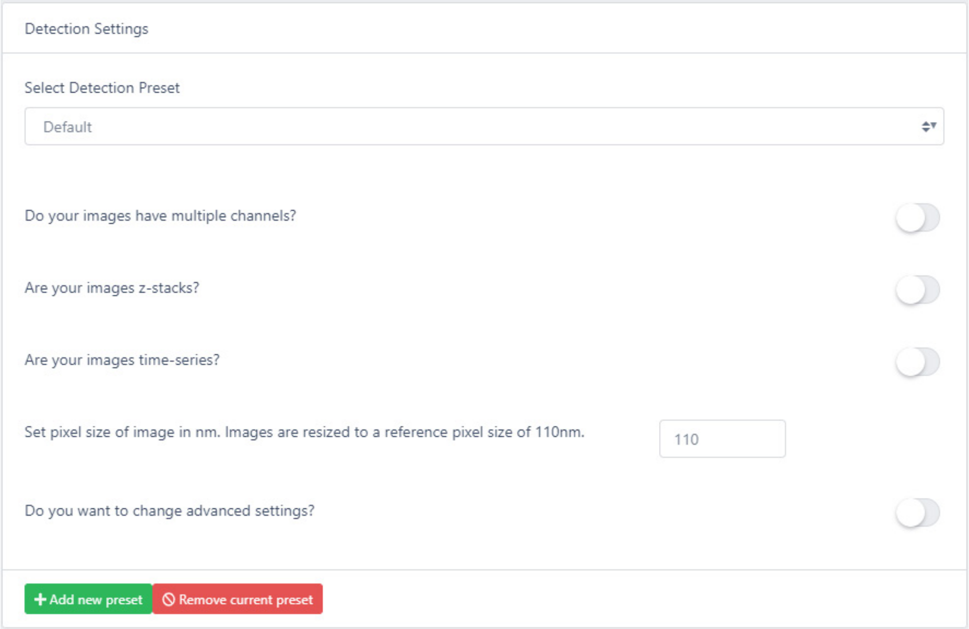

as below:

You may change the default pixel size according to your specific image parameters, but if it’s not significantly different, YeastMate will work well.

3) Back in the ‘Start new job’ page, press the ‘Start backends’ Button. Two console windows will open. Both IO Backend and Detection Backend markers will turn green.

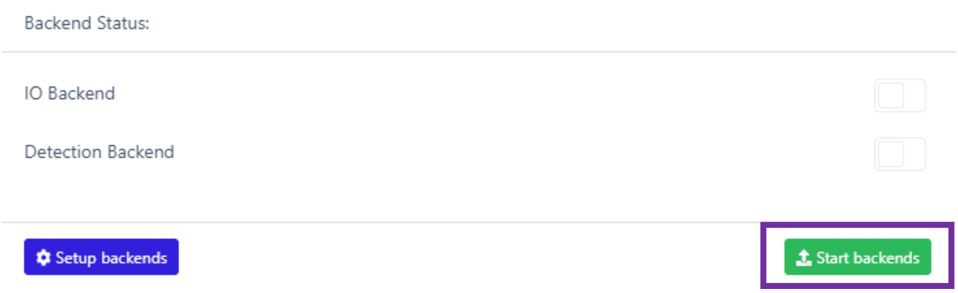

**Script Steps:**

1) First, the user is prompted to choose the folder with the cell images in visible light (*i.e.* brightfield or DIC).
2) The user is asked to make sure they opened YeastMate. After making sure YeastMate is ready (*i.e.* backends are running), and clicking ‘OK’, the script will continue.

3) For each image in the folder:
   I. The image is opened.
   II. For each slice: YeastMate will create a multi-colored mask image. Masks will be transformed to binary images.
   III. Using Analyze → Analyze Particles, cells are added to the Roi Manager and counted.
4) The plane with the largest number of identified cells is chosen.
5) The chosen plane is duplicated: Image → Duplicate; *n*=number of original planes.
6) The images are converted to binary images: Process → Binary → Convert to Mask.
7) A new, stacked binary image is created: Image → Stacks → Images to Stack.
8) The image is saved in the same folder as the original images, under the 'Cells Masks' folder.

For each image you will get a binary image, with a white background and the shape of the cells in black. This will be used later to identify each cell's location.

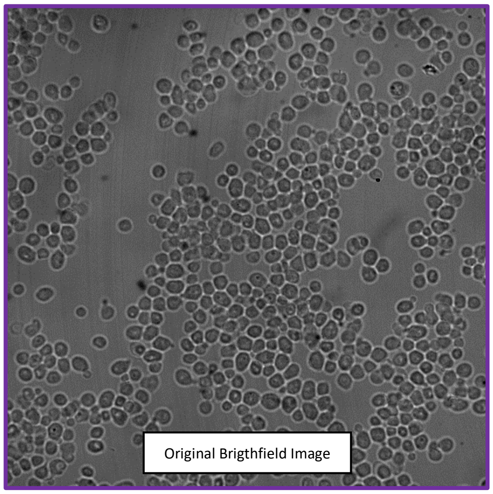

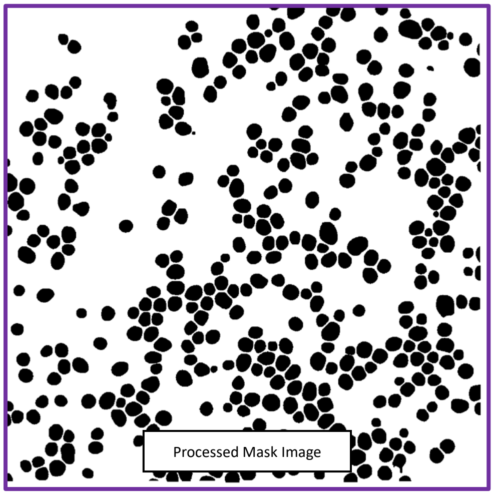

**Unsolved Bugs or Issues:**

Low quality images, where no cells are identified may occasionally lead to YeastMate crashing for one or more of the *z* planes, resulting in the script crashing as well. In this case, the image cannot be used and must be removed from all folders (*i.e.* all channel images) before running the script again. Also, before re-running the script, make sure to delete all the folders added in the disrupted run, to avoid any future crashes. Once the script is done, the Fiji ‘Console’ window may open, presenting an error. It can be ignored.

#### 5. Use Trainable Weka Segmentation to Train a Model for Sub-cellular Segmentation

Identifying and defining sub-cellular structures is a more challenging step, especially when the structure is irregular. For example, the shape of the endoplasmic reticulum (ER) varies considerably from cell to cell. The solution from Weka, in the form of a Fiji plugin, allows a user to easily train a model and use the results of the training (*i.e.* a file called a classifier or a model) to perform segmentations of all similar signals in the different images. Here follows a detailed explanation of the training process.

**Classifier Training Steps:**

1) Open an image of the appropriate channel in Fiji.
2) If you wish to save time, make sure to only use a single *z* plane image for this stage (in a multiple stacks image: Image → Stacks → Stacks to images and select a *z* plane window with a good signal).
3) Press Plugins → Segmentation → Trainable Weka Segmentation. The pictured window will open (Image 1).

1

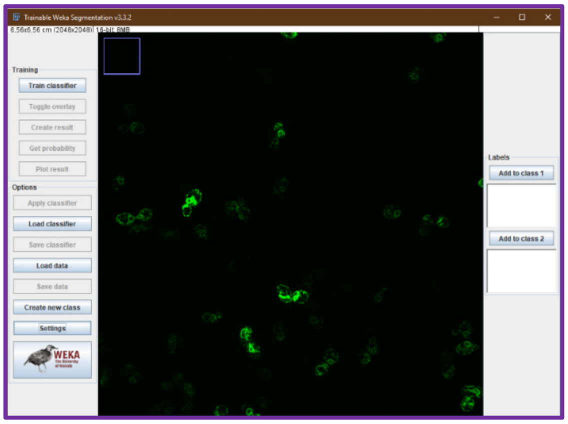

4) To train the classifier, you must mark specific pixels of each class (*e.g.* your organelle, background etc.). Name each class by pressing ‘Settings’, at the bottom left (Image 2).

2

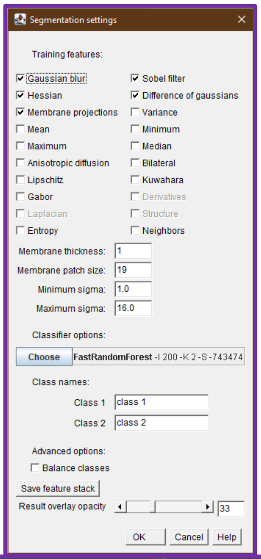

5) Change class names to fit your experiment (for example, ‘ER’ and ‘Background’).

6) Under the settings window, you may also change different features that can alter the behavior of the classifier. Notice adding more training features will result in longer training and identification times, and require more computing resources, but feel free to experiment. The settings in the picture are the ones used by us.

7) Now you can mark different classes on the image using the mouse. It is recommended to adjust brightness/saturation and zoom-in for this stage (note: adjusting brightness won’t affect the classifier).

4

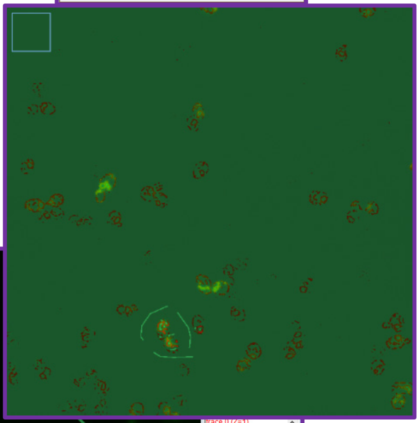

8) Mark a small area of the image each time, and press ‘Add to Class’ button. The markings will change color and be added to the list under the class name (Image 3).

3

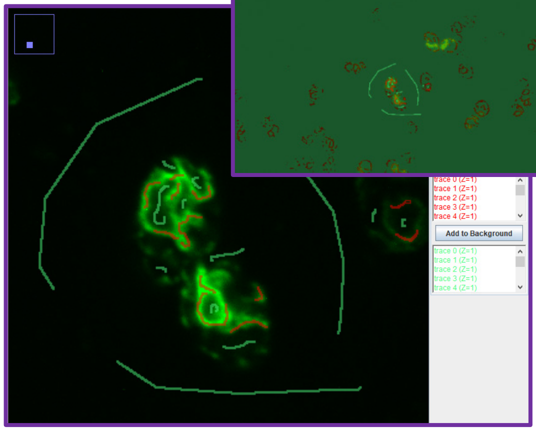

9) If the marking is inaccurate, avoid adding it to a class. You cannot remove a marking once it’s added. You can start marking without saving and the previous marking will disappear.

10) After marking (and adding each) a few times for each class (no need to add more than 3-4), press ‘Train classifier’.

11) After training ends, you will be able to see how the different classes are marked on the entire image (Image 4).

12) Save the classifier by pressing ‘Save classifier’.

13) Repeat these steps for several different images (use the 'Load classifier' button and select the classifier you created). The more images you use, the better, but only a few marks in each image are enough.
14) You can test and re-train your classifier on different images of the same channel/signal. By pressing 'Get probability', you can look at a probability map resulting from your classifier. Make sure to look at it next to the original image and make sure the signals match. If they don't match, you need to retrain your classifier.

#### 6. Perform Sub-cellular Segmentation Using a Trained Model

The following script uses a trained classifier on multiple images in a user input folder. The script assumes a classifier was already created (as described above), and the input folder contains only images of a channel corresponding with the chosen classifier.

The script saves both the organelle mask and statistical map files. The statistical maps are not used downstream by other scripts but can be important for troubleshooting. After going over a few of them to make sure organelle segmentation was performed well, they can be deleted.

Note that this step is very time-consuming. When using a classifier for the first time, it is recommended to test it by using the script on a few images (*e.g.* 3-4) and examine the resulting masks to make sure the results are of high quality.

**Script Steps:**

1) The user is asked to input the folder containing images of the channel they wish to segment and the classifier (through a browser window).
2) Subfolders are created to save the probability maps and mask images in the input folder.
3) For each image:
    I. Each *z* plane is saved as a separate file (*i.e.* in a temporary folder: 'separated_planes').
    II. Every separate *z* plane image is opened one at a time and the following actions are performed:
        i. Trainable Weka Segmentation is activated.
        ii. Using 'Load classifier' the inputted classifier is loaded.

iii. Using ‘Create a probability map’ produces the probability map.
iv. Probability map is saved in a subfolder (in a temporary folder: ‘temp_maps’).
v. Using ‘Close all’ to close all image windows.

III. Once all planes from one image have a probability map, the maps are opened.

IV. All *z* planes are joined to a single image using ‘Stack to a single image’ (under the same name + ‘_map’).

V. The probability map image with multiple *z* planes is saved in the probability maps folder.

VI. A mask file based on the probability maps is created (using IsoData thresholding) and saved.

VII. All image windows are closed using ‘Close all’.

VIII. All temporary image files from ‘separated_planes’ and ‘temp_maps’ folders are deleted.

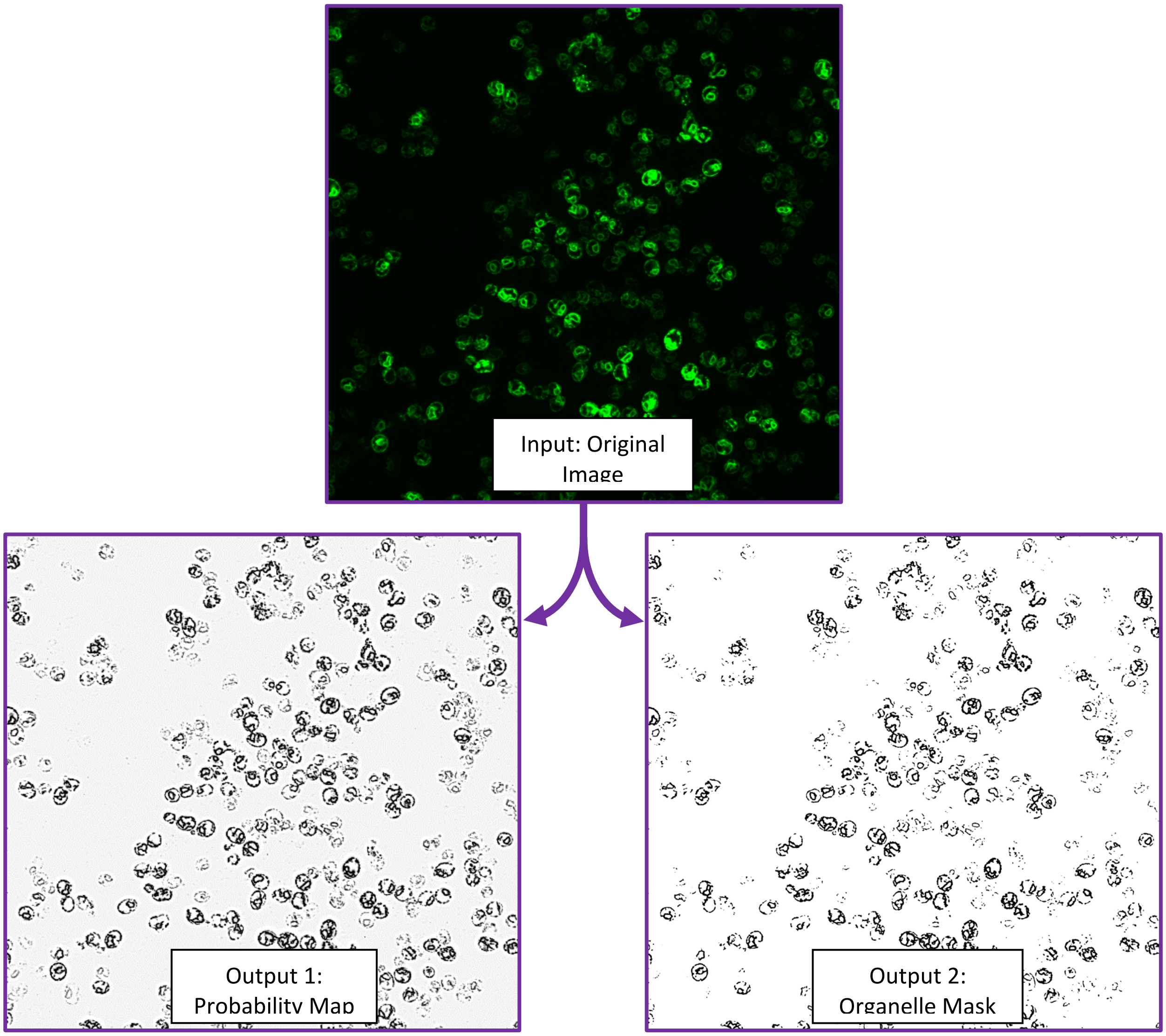

#### 7. Organelle Sub-segmentation

The ER can be divided into cortical (cER) and perinuclear (nER) sub-sections. The latter is proximal to the nuclear membrane and the former extends from the nucleus and localizes distally (*e.g.* under the plasma membrane). When checking mRNA colocalization with the ER, it can be calculated separately per sub-section. EASI-ORC can separate the two sub-sections by using an image of the nucleus as a proximity marker. The nucleus is a specific case of a proximity marker that allows for the sub-segmentation of an organelle. The process can be performed with any proximity marker you choose in order to sub-segment previously made masks.

This script assumes the Trainable Weka Segmentation process was performed on both organelle and proximity marker signals and masks exist for each. Each mask image must be saved in their own folder (without any subfolders) and the files must be ordered in the same manner. If the files were created using the scripts in this guide, all these conditions are met.

**Important note:** In cells or *z* planes lacking a proximity marker signal, there is no way of identifying the sub-sections of the organelle that are either proximal or distal to the marker. As such, these cells in these planes must be filtered out in the colocalization calculations (the EASI-ORC colocalization script takes this into account).

**Script Steps:**

1) The user is asked for the folder of the proximity marker and organelle masks.
2) The user is asked for the number of dilation steps (*i.e.* adding 1 row of pixels near signal edges) to be performed on the proximity marker signal.

   ***Note:** It is recommended the user tests the number of dilations manually with several values, to make sure the proximity marker is of satisfactory size to cover the proximal sub-section (*e.g.* for ER sub-segmentation using a nuclear marker, we found that 3 steps works well). To check for yourself, simply open a proximity marker mask and go to the menu: Process → Binary → Dilate. You can then merge the proximity marker image with the organelle image and see if the coverage is sufficient.
3) For each proximity marker image:
   I. Process → Image Calculator… This performs subtraction of the proximity marker mask image from the organelle mask image, which outputs a mask image containing only the distal sub-section.
   II. Process → Image Calculator… This performs subtraction of the produced distal sub-section mask image form the original organelle image, which outputs a mask image containing only the proximal sub-section.

All images are saved in the original organelle mask folder. The process and the effect of different qualities of proximal markers are shown below. Panel C exemplifies proximity marker quality by showing nER and cER sub-segmentation in cells with differently marked nuclei.

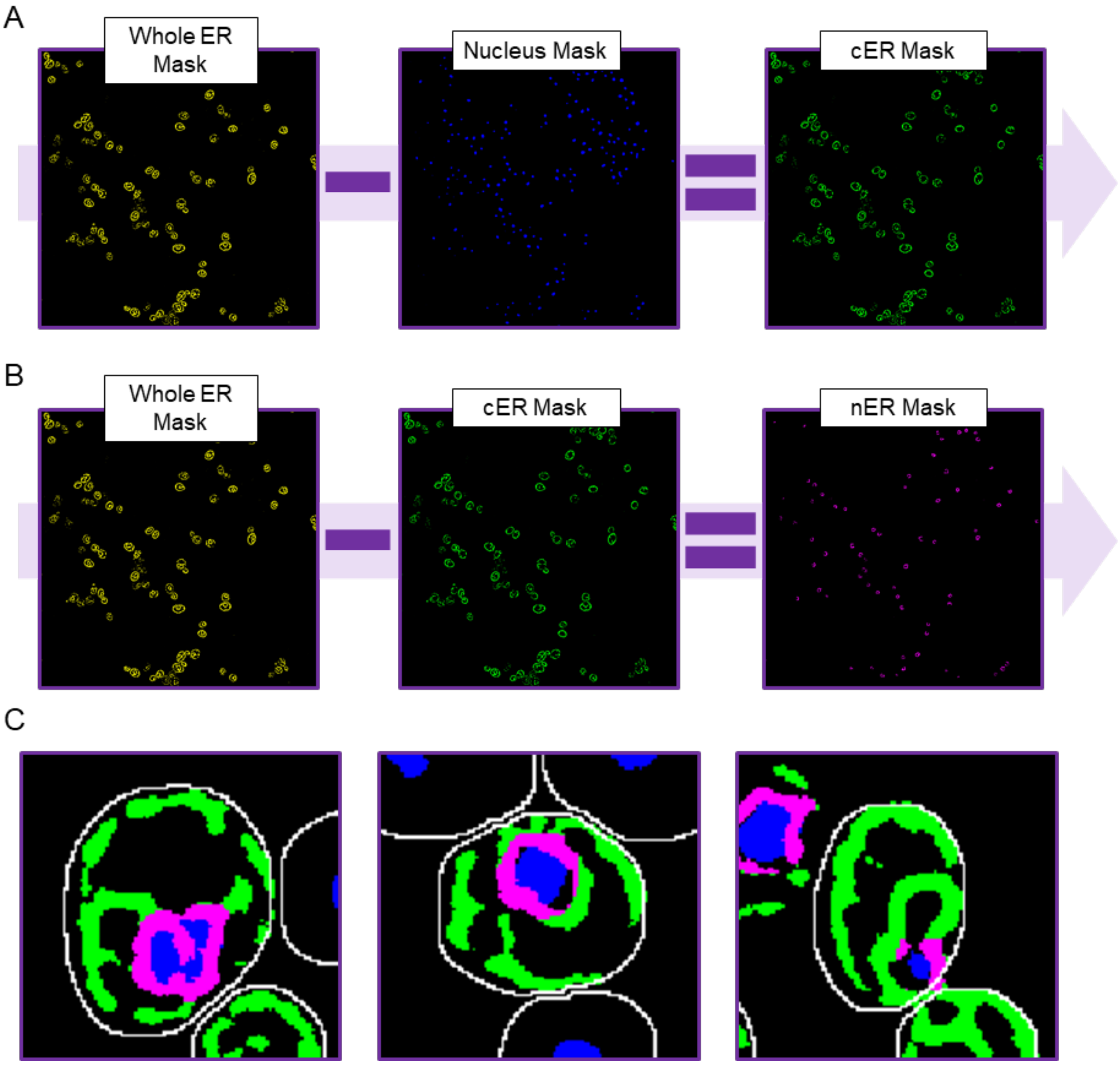

#### 8. Manual run of RS-FISH for threshold identification

smFISH signals are relatively simple signals that can be identified using specific plugins. RS-FISH is one such plugin. It's fast, accurate, and considers the signal intensity between the different *z* planes. Crucially, it can be automated using Fiji scripts.

**Important:** To properly identify FISH spots, RS-FISH requires a signal threshold which will be different between experimental procedures and different probe sets. So before automation, RS-FISH must be used manually on a few images for each signal type you wish to identify. A step by step explanation is provided here, but to gain a full understanding of the process, it is recommended to read the documentation provided with the plugin. Make sure to follow the link provided in the Introduction to install RS-FISH before continuing.

**Threshold Value Identification:**

1) Open an mRNA channel image and activate the plugin (Plugins → RS-FISH → RS-FISH).
2) A window will open. Under 'Mode', choose 'Interactive'. Under 'Robust fitting', choose RANSAC. Tick both boxes and under 'Spot intensity', choose 'Linear Interpolation' (it should look like it does in Image 1). Click ok.

1

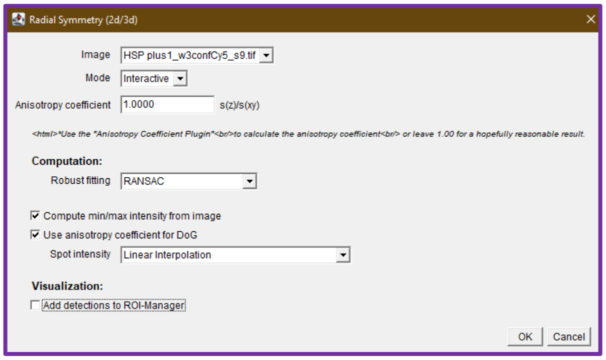

3) Several windows will pop up. Move the additional image window side-by-side with the original image.
4) If you haven't done so before, change the image's brightness and contrast so you can clearly see signals. Drag and move the square that appeared in your image, so it'll mark an area with signals contained in it. You can also manipulate its size and shape of the area to your liking, though it shouldn't be larger than a couple of yeast cells. Zoom in to it on both images.
5) Now, when you move across different slices, you can see signals are marked by a red circle. For each spot across all planes, only the most intense are marked.
6) In the 'Adjust difference-of-gaussian values' window (Image 2), you can change the Threshold value. Change it several times and test to see the best value that allows you to

2

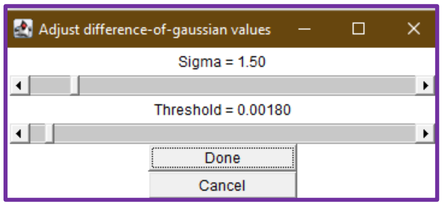

capture as many signals as possible, without marking noise. It is always preferable to miss some signals (*i.e.* false negatives) than to mark noise as a signal (*i.e.* false positives).

7) Once an appropriate value is found, you may check it by repeating the process on a second and third image with the same signal source.
8) The threshold value can be used on all images created using the same signal source (*i.e.* a specific probe or other marker) that were subjected to the same capturing method (*i.e.* same channels, same microscope, same exposure time, etc.).

#### 9. smFISH spot identification using RS-FISH

Once you have an appropriate threshold for your FISH signals, you can use the automation script to collect data regarding the location of all spots in your images.

**Script Steps:**

1) The user is asked for the location of the smFISH channel images directory.
2) The user is asked for the spots identification threshold value.
3) Once these are inputted, the script performs the following steps on each image:
    I. Opens smFISH image
    II. Runs RS-FISH, using the provided spots threshold
    III. Sorts results by '*z*' value (the *z* plane where it was found)
    IV. Saves results using current image name

Result tables will be saved as csv files, under the same directory as the smFISH channel images, in a new directory called 'smFISH Spots'. The output table for each image will include data on each point (from left to right): *x*, *y*, *z* coordinates (width, height and *z*-plane, respectively), time (only relevant for time lapse images), channel (not relevant with single channel images) and intensity (see table below).

| x | y | z | t | c | intensity |
|---|---|---|---|---|---|
| 1519.123 | 65.0544 | 1.1473 | 1 | 1 | 252.9637 |
| 1203.276 | 48.6669 | 1.1954 | 1 | 1 | 120.9883 |
| 1588.631 | 85.913 | 1.2808 | 1 | 1 | 357.8814 |
| 211.1326 | 1167.16 | 1.3055 | 1 | 1 | 246.1136 |
| 582.0259 | 364.5312 | 1.4322 | 1 | 1 | 221.7589 |

#### 10. Colocalization of smFISH signals and organelle

Once the organelle is defined in the mask files and smFISH spot data is obtained via RS-FISH, colocalization can be examined. This script is the most complex one, so a general description of its steps is provided, instead of a step-by-step account.

First, this script will prompt the user for the locations of the cell mask images and smFISH spot csv files created by RS-FISH (if you wish to identify spots using another method, provide a csv file with the same structure as the RS-FISH output as presented above).

Next, the script will ask the user whether the organelle was sub-segmented (as described in Section 5 of this guide). If it was, choose 'Yes'. If it wasn't, choose 'No'. According to the user's answer, they will be prompted to provide locations of organelle and proximity marker masks (*i.e.* only if organelle sub-segmentation was performed) and a minimum value for the organelle coverage (*e.g.* for nucleus, we used 0.05 for 5% cell coverage). Make sure you input the binary mask file directories and not the statistical map file locations.

The user will be asked where they want to save the Results tables and lastly, for the value of the smFISH signal diameter, in pixels. We recommend using 1, as this will only look for signals 1 pixel away from the spot. If you wish to choose a different value, you can examine several images closely, by zooming into several isolated signals and count the number of pixels marking their diameter. Colocalization is measured according to the diameter chosen. A large diameter may mark signals far from the organelle as "colocalized", and a small one will result in less signals identified as such. Take that into consideration when choosing this value. Make sure to use the same signal diameter when examining the same signal source (*i.e.* the same FISH probe set and target, for example).

After the user inputs the values and folders, the script goes over each image, each plane within that image, each cell, and each mRNA. Each mRNA is matched with its specific cell, as well as given a colocalization assignment to the organelle. Colocalization is defined by having **more than 0** organellar signals contained within the diameter provided by the user. In case both sub-section signals are present, the one with a greater total signal (*i.e*. having more organelle signal pixels) is chosen. In case of a tie between the signals, the check will be repeated sequentially, with bigger diameters (starting from 1), until the tie is broken. The check is done by creating a circle with the provided radius around the smFISH spot's maxima and measuring the amount of organelle signal inside that circle (using basic ImageJ commands). Note that since RS-FISH uses data from multiple slices to identify a spot's location, its '*z*' value can be smaller than 0 or larger than the actual number of cross sections in the image (meaning that it is located at a theoretical point, beyond the limits of the image). In these cases, the spot will be ignored since there is no organelle data for that location. The number of spots of this nature, if any, is very small. In addition, the *z* plane chosen for colocalization measurement will be the nearest one to each spot. The script filters out cells according to a few features*, e.g.* edge cells (containing pixels found in the 2% nearest to the edges) and cells with less than the input cellular area of proximity marker signal, per plane.

For each image, a CSV file is created in the folder the user has inputted. The results will be saved in the following format: Cell #, total number of mRNAs in cell, number of mRNAs of each localization type (organelle proximal/distal, non-organelle), a list for each spot within that cell (Each spot data appears in this order: X,Y coordinates, *z* plane, spot intensity, localization determined) and the organelle coverage value for each z-plane in the image (see table below).

| **Cell #** | **Total mRNA per Cell** | **Total Colocalized With Organelle Proximal** | **Total Colocalized With Organelle Distal** | **Total Not Colocalized with Organelle** | **Spots Coordinates Intensity and Colocalization (Distal Proximal or Not Colocalized)** | **Organelle Signal Size Cross Section 1** | **Organelle Signal Size Cross Section n** |
|---|---|---|---|---|---|---|---|
| 12 | 3 | 0 | 1 | 2 | [[349.8452,131.7305,11,155.3453,organelle_distal],<br>[312.4651,136.6522,13,161.7325,nc],<br>[329.2508,163.6214,13,147.0048,nc],] | 24.586 | 37.651 |
| 31 | 1 | 1 | 0 | 0 | [[1411.1692,295.842,5,174.1844,organelle_proximal],] | 56.234 | 43.354 |
| 34 | 1 | 0 | 0 | 1 | [[1376.327,323.9262,13,202.1867,nc],] | 0.110 | 1.397 |

These files can be analyzed by any means you wish, but we recommend the use of our analysis toolset. A user guide for it can be found in the next section.

**Important:**

The script must run on folders containing only the images described above and no other file or folder. Using the previous scripts in the guide to create these images will make sure all files are saved and ordered appropriately. Lastly, the script cannot run in batch mode. Meaning, it must open each image it uses and should not be interrupted while it works. **It is recommended you do not use the computer while it is running** (it's quick, but if you have many images, try to run it on 2-3 of them to evaluate how long you need to leave your computer. For example, ~50 images shouldn't take more than 15 minutes).

**Unresolved Bugs or issues:**

At times, an empty results table window remains open after the script is done. It can be ignored and closed.

The log will be filled with lines only containing '1'. This is an output of the file deletion function and can be ignored.

#### 11. Colocalization data analysis tool

To allow easy analysis of the EASI-ORC output files, we have developed and provided several tools written in Python that can be used either locally or online, via Jupyter Notebook or Google Colab, respectively (the local version requires Python installation, while the online version does not). The notebooks also include directions for each step before each code cell. There's no need for coding in order to use the analysis tool. Only pressing the 'run' button of each function you wish to activate (*e.g.* filtering, graphing, and statistical analysis output). However, if used locally, you will need to install Python and several packages.

Here, we explain what the analysis tools provide. Specific instructions for each section can be found in the analysis tool notebook itself.

The analysis tool can be divided into **four sections**:

1) **Table consolidation.**

2) **Data Filtering.**
3) **Data Visualization.**
4) **Statistical Analysis.**

Section 1 asks the user how many different treatments/strains are present in the experiment. In the online version, it will prompt the user to select the files for each treatment and name each one (this name will appear in Results tables and graphs). In the local version, the user is asked for the folder that contains the results, identification strings of each sample (which must be specific and are case-sensitive), and the name the user gives to each sample to be displayed in the tables and graphs. It will then create a consolidated table for each treatment, which can be used in the next sections. The consolidated table has each FISH signal represented by a line, with data points for *x*, *y* coordinates, *z* slice, intensity, cell id, colocalization designation (1 if it belongs to the designation an 0 if not), corresponding organelle coverage and treatment.

Table 1: Example Consolidated Data Table

| Data Point | Value | | |
|---|---|---|---|
| x | 1315.1729 | 1034.355 | 1080.607 |
| y | 466.532 | 1630.217 | 311.2029 |
| z | 2 | 7 | 4 |
| intensity | 339.3091 | 304.3342 | 278.8375 |
| cell_id | 6 | 676 | 9 |
| Colocalized | 1 | 0 | 0 |
| not_colocalized | 0 | 1 | 1 |
| organelle_proximal_colocalized | 0 | 0 | 0 |
| organelle_distal_colocalized | 1 | 0 | 0 |
| organelle_coverage | 28.01095 | 22.05703 | 35.62368 |
| Treatment | Strain 1 | Strain 2 | Strain 3 |

Section 2 provides several filtering steps that are presented visually (*i.e.* via histograms that display the data after filtering) and updated as you input new filtering values. The filters must be performed in the order in which they appear (*i.e.* if you change one, you must also change the next ones). Each filter has minimum and maximum values. First, there is a filter for smFISH spots by spot intensity.

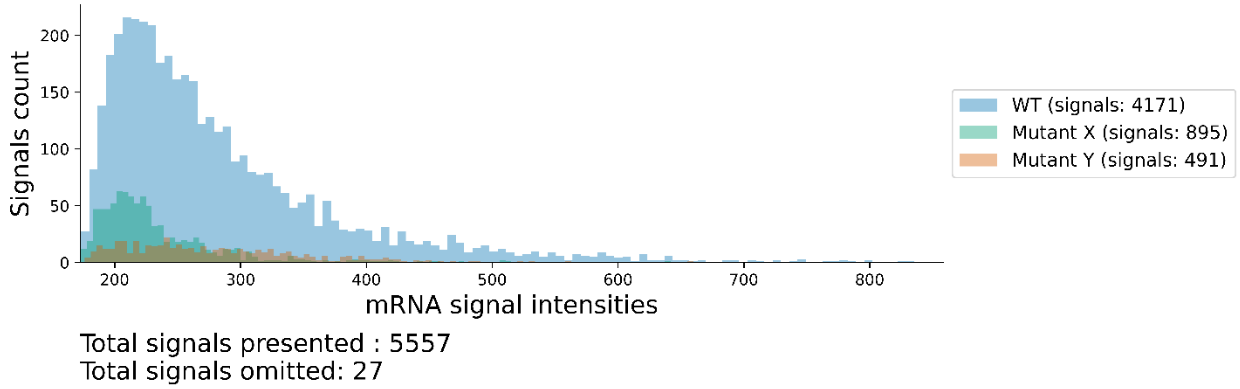

**Filter 1 – Intensity per mRNA signal**

Second, there is a filter to remove *z* planes according to organellar signal coverage per cell and per *z* plane (note that a organellar signal of zero will lead to a non-colocalized designation to all FISH spots, while too high signals will likely designate all FISH spots as colocalized).

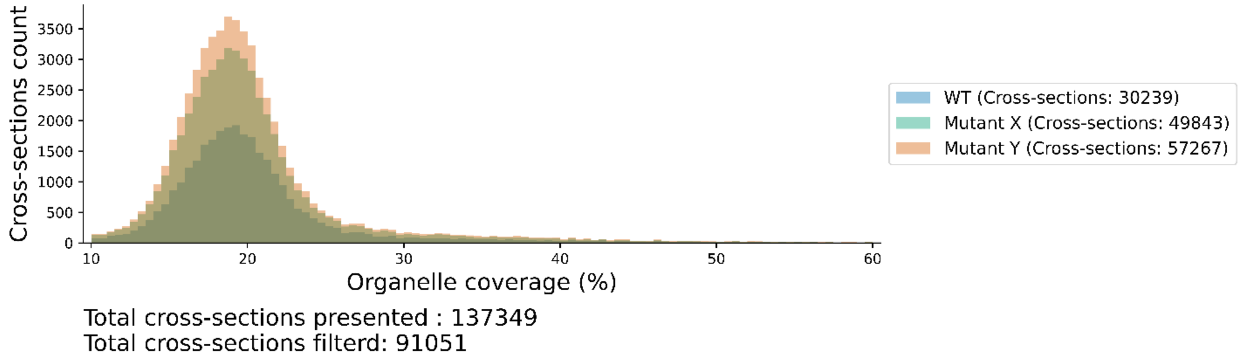

**Filter 2 – Organelle coverage of cell per cross-section**

Third, you may filter the data by the number of FISH signals scored per cell.

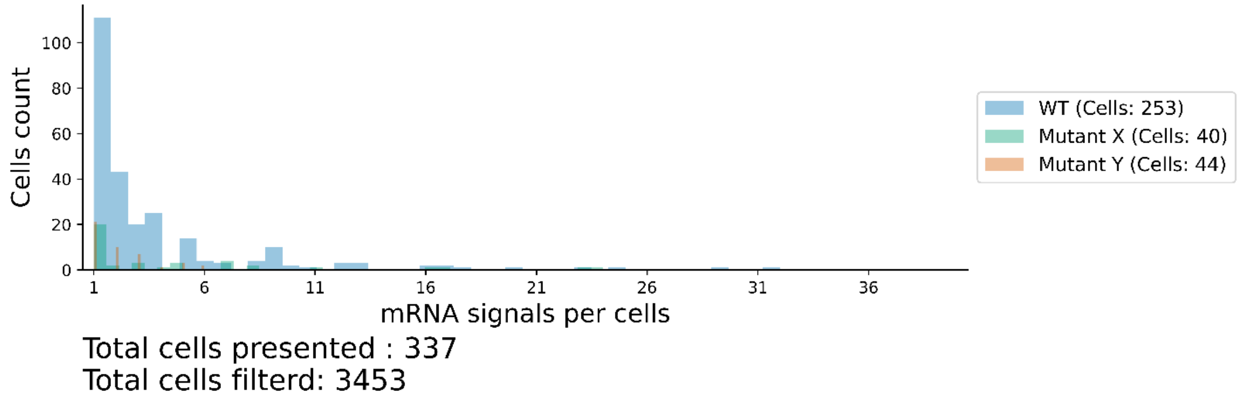

**Filter 3 – Number of mRNA signals per cell**

This section also creates a new table for each treatment of the data with the filters applied to it. Filter values can be inputted directly (by double clicking the number and changing it) or using the sliders above the graph.

Section 3 provides several graphs for colocalization analysis: A proportion graph representing the percentage of mRNA-organelle localization (or mRNA colocalization with the proximal and distal organelle sub-segments, if sub-segmentation was performed), violin graphs of each colocalized sub-population (*i.e.* proximal, distal, non-colocalized and total colocalized proportions), and violin graphs showing the number of mRNA signals per cell, per treatment, and average organelle coverage (calculated only from the *z* planes remaining after filtering for organelle coverage).

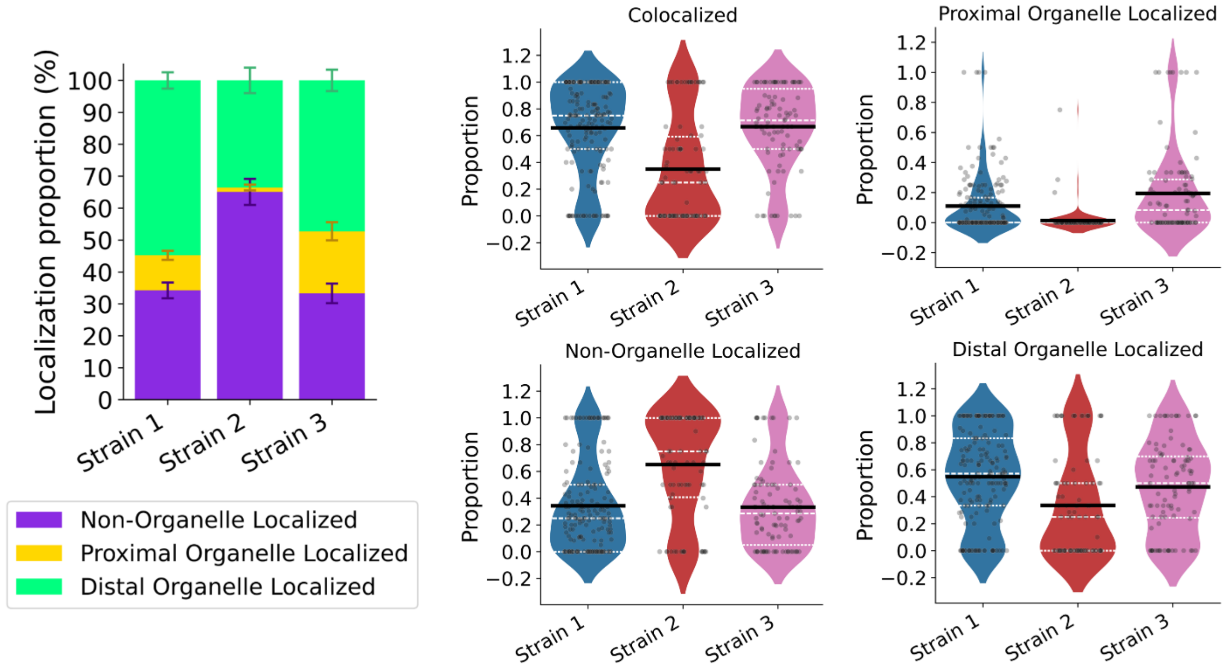

**Output Plots: mRNA-organelle Colocalization**

In addition, two scatter plots are produced to describe the correlation between mRNA-organelle colocalization proportion with either the number of mRNA signals per cell or the average organelle cellular coverage (the average is calculated between all *z* planes scored).

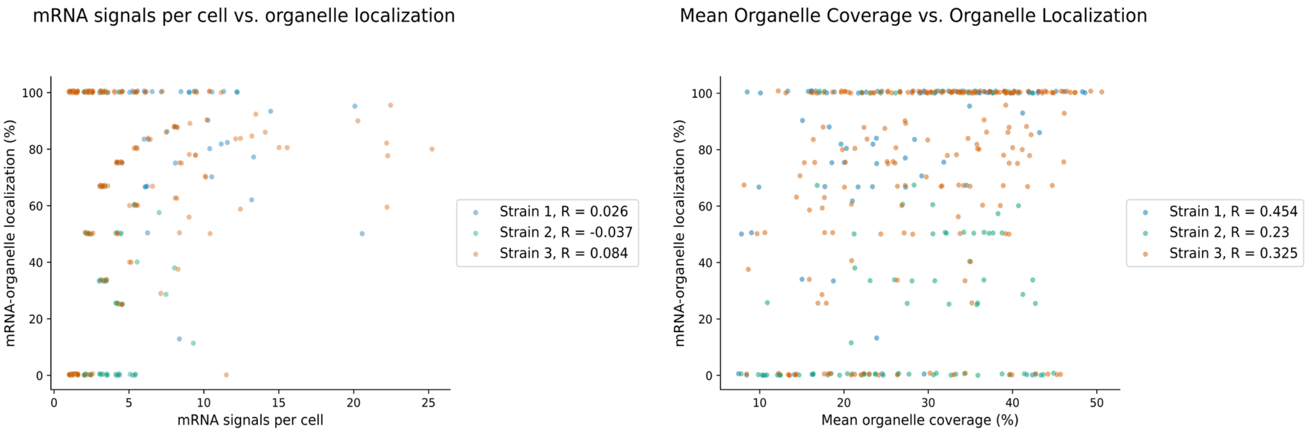

**mRNA-organelle Colocalization Correlation plots**

Finally, Section 4 provides a table with statistical tests performed for and between each treatment. Each treatment's data set is compared using an ANOVA test to every other treatment, to provide statistical significance of the differences between each treatment. All results are collected in a statistical table, saved as a csv file (Table 2). A Tukey's test is also performed and saved in a separate file.

Every graph and table produced by the analysis tool is available to be saved individually and downloaded together, in a zip file in the online version.

Table 2: Example of Statistical Summary Table

| Feature | **Strain 1** | **Strain 2** | **Strain 3** |
|---|---|---|---|
| Number of cells before filtering | 1164 | 690 | 785 |
| Number of cells after filtering | 180 | 90 | 100 |
| Percentage of positive cells | 15.4639 | 13.0435 | 12.7389 |
| mean proportion of organelle localization | 0.6602 | 0.3492 | 0.6624 |
| standard deviation proportion of organelle localization | 0.3307 | 0.3871 | 0.3170 |
| standard error proportion of organelle localization | 0.0246 | 0.0408 | 0.0317 |
| mean proportion of non-organelle localization | 0.3398 | 0.6508 | 0.3376 |
| standard deviation proportion of non-organelle localization | 0.3307 | 0.3871 | 0.3170 |
| standard error proportion of non-organelle localization | 0.0246 | 0.0408 | 0.0317 |
| mean proportion of organelle proximal localization | 0.1092 | 0.0137 | 0.2031 |
| standard deviation proportion of organelle proximal localization | 0.1915 | 0.0866 | 0.2939 |
| standard error proportion of organelle proximal localization | 0.0143 | 0.0091 | 0.0294 |
| mean proportion of organelle distal localization | 0.5511 | 0.3355 | 0.4593 |
| standard deviation proportion of organelle distal localization | 0.3397 | 0.3792 | 0.3351 |
| standard error proportion of organelle distal localization | 0.0253 | 0.0400 | 0.0335 |
| mean mRNA signals per cell | 9.7056 | 2.7556 | 10.5900 |
| standard deviation mRNA signals per cell | 10.2960 | 1.7883 | 12.7439 |
| standard error mRNA signals per cell | 0.7674 | 0.1885 | 1.2744 |
| mean organelle coverage | 25.9877 | 28.0908 | 25.4835 |
| standard deviation organelle coverage | 10.2517 | 9.9213 | 9.0045 |
| standard error organelle coverage | 0.7641 | 1.0458 | 0.9005 |
| Pearson correlation mean organelle coverages vs organelle localizations | 0.3293 | 0.2332 | 0.4323 |
| Correlation p value mean organelle coverages vs organelle localizations | 6.38E-06 | 0.0270 | 7.08E-06 |
| Pearson correlation mean mRNA signals vs organelle localizations | 0.1059 | -0.0424 | 0.0612 |
| Correlation p value mean mRNA signals vs organelle localizations | 0.1570 | 0.6914 | 0.5452 |
| ANOVA (one-way) colocalized p value | 3.57E-12 | 3.57E-12 | 3.57E-12 |
| ANOVA (one-way) colocalized statistic | 28.3466 | 28.3466 | 28.3466 |
| ANOVA (one-way) not_colocalized p value | 3.57E-12 | 3.57E-12 | 3.57E-12 |
| ANOVA (one-way) not_colocalized statistic | 28.3466 | 28.3466 | 28.3466 |
| ANOVA (one-way) organelle_proximal_colocalized p value | 6.97E-09 | 6.97E-09 | 6.97E-09 |
| ANOVA (one-way) organelle_proximal_colocalized statistic | 19.7771 | 19.7771 | 19.7771 |
| ANOVA (one-way) organelle_distal_colocalized p value | 1.31E-05 | 1.31E-05 | 1.31E-05 |
| ANOVA (one-way) organelle_distal_colocalized statistic | 11.5951 | 11.5951 | 11.5951 |
| ANOVA (one-way) rna per cell p value | 1.50E-08 | 1.50E-08 | 1.50E-08 |
| ANOVA (one-way) rna per cell statistic | 18.932 | 18.932 | 18.932 |
| ANOVA (one-way) filtered_mean_organelle_coverage p value | 0.149 | 0.149 | 0.149 |
| ANOVA (one-way) filtered_mean_organelle_coverage statistic | 1.914 | 1.914 | 1.914 |
